# Volumetric printed biomimetic scaffolds support *in vitro* lactation of human milk-derived mammary epithelial cells

**DOI:** 10.1101/2025.03.18.643642

**Authors:** Amelia Hasenauer, Kajetana Bevc, Maxwell C. McCabe, Parth Chansoria, Anthony J. Saviola, Kirk C. Hansen, Karen L. Christman, Marcy Zenobi-Wong

## Abstract

The human breast is remarkably plastic and remodels with each birth to produce milk optimally suited for the changing demands of the newborn. This dynamic nature of lactation makes it challenging to study under controlled conditions. Given the health benefits of human milk, models of secretory mammary tissue would offer new opportunities to study factors that influence this important food source. First, 3D models of the mammary duct/alveoli (D/A) were designed based on shapes found *in vivo*. Photoresins based on mammary decellularized extracellular matrix (dECM) were optimized to match the mechanical properties of native breast tissue. Next, these dECM-based D/A models were printed with a volumetric printer and seeded with human milk-derived mammary epithelial cells (MECs). MECs formed stable epithelial layers on the printed surfaces and secreted the beta-casein and milk-fat-globules. This novel model offers exciting avenues to explore hormonal, nutritional, and mechanobiological factors involved in lactation, thereby improving understanding of lactation for the benefit of infant and their mothers.

## Introduction

Human breast milk bolsters the immunity and microbiome health of newborns, assists neuro-cognitive development, and is critical to the prevention of multiple debilitating childhood diseases (1, 2). Unfortunately, only 30-60% of infants are fully or partially breastfed during the first six months after birth, and alternatives (i.e., bovine milk-based infant formula) lack many of the constituents associated with health benefits (3, 4). Accurate models of human lactation would allow us to address the fundamentals of lactation biology and identify environmental factors that influence milk production and composition.

Mammary epithelial cells (MECs) are the milk-producing cells of the mammary gland. Luminal MECs synthesize milk components supported by basal MECs that contract to facilitate milk secretion (5–7). Bissell and coworkers have fundamentally reshaped our understanding of the extracellular matrix (ECM) and emphasize its role beyond a mere passive support scaffold to an active regulator of cell signaling and behavior (8, 9). MECs interact with the basement membrane and underlying ECM through integrin-mediated binding and respond to mechanical and biochemical stimuli of the ECM, to induce cell polarization along the basal/apical axis, tight junction formation, and tissue self-reorganization (5–7).

Despite recent advances in tissue engineering, there are very few models of *in vitro* lactation and those that exist rely heavily on organoids. Organoid technology is a versatile tool that allows one to explore the mammary gland *in vitro* from development to breast cancer. However, organoids face limitations such as low formation rates, limited life span, and variations in shape and/or size that contribute to experimental variability. Furthermore, their use precludes the continuous collection and analysis of secreted milk proteins without cell layer disruption. Most *in vitro* models also rely on murine cells, which vary in developmental timing, cell diversity, and ECM composition compared to humans (11, 12). Additionally, murine milk differs from human milk in cell, protein, fat, and carbohydrate content reflecting species-specific nutritional needs (13, 14). These differences can compromise the translational relevance of organoid work, which further underlines the need for alternative human breast models.

To propose a new approach, we used light-based 3D printing to biofabricate human lactation models composed of decellularized ECM from bovine breast tissue and human milk-derived mammary epithelial cells (MECs) (Fig. 1).

**Figure 1:**
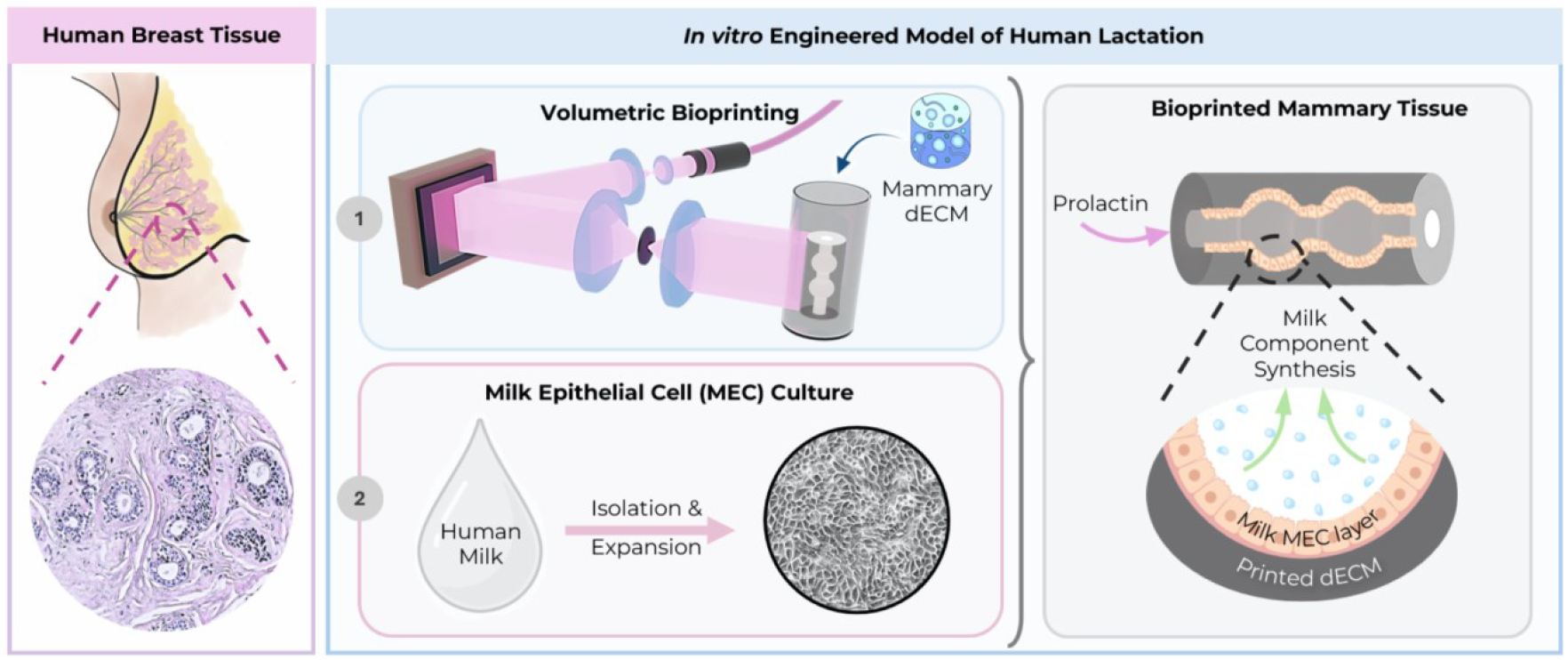
3D printing of human alveolar tissue. Left: In human breast tissue, alveoli are the milk-producing units during lactation. Right: (1) Volumetrically printed (VP) decellularized extracellular matrix (dECM) and (2) milk-derived mammary epithelial cells (MECs) are leveraged to create secretory in vitro models. Expanded milk MECs are post seeded in the printed mammary tissue structures where they synthesize milk.

To date, the literature reports only a few studies that involve bioprinting of MECs, primarily focused on breast cancer organoids combined with extrusion bioprinting (15, 16). In these studies the placement and number of cells (or organoids) within the biomaterial can be precisely controlled, which reduces experimental variability (17, 18). However, current extrusion-based approaches have not successfully replicated the architecture of mammary ducts and alveoli, primarily due to challenges associated with printing overhanging hollow shapes. Thus, current studies rely on the cells’ ability to self-organize into these structures, which restricts the spatial control over the epithelial and stromal compartment’s architecture, particularly given the inherent biological variability in tissue formation (16).

In contrast, recent advancements in volumetric printing (VP) overcome the limitations of conventional layer-by-layer additive manufacturing and allow for the printing of perfusable tissue structures. In VP the 3D model is discretized into a series of images which are projected onto a rotating vat filled with photoresin. The voxels where the accumulation of photons surpasses the gelation threshold undergo solidification, while the remaining resin can be washed away. This cutting-edge technology marries high resolution with fast printing speeds (several seconds) to produce defect-free structures with precisely controllable surface topographies, which makes it an attractive yet unexploited choice for engineering mammary tissue (19, 20).

The recent progress in hardware printing has been complemented by advances in biomaterials. One such innovation is tissue decellularization, which transforms organs and tissues into cutting-edge biomaterials that can be directly light-printed using VP without any chemical modification (21, 22). The interest in decellularized extracellular matrix materials (dECM) stems from the preservation of soluble proteins during decellularization, which enhances the extracted biomaterials’ bioactivity and reduces cytotoxicity. This makes dECM-based resins more conducive to cell-matrix, proliferation, and differentiation compared to single component resins like methacrylated biomaterials (e.g., Gel-MA) and ultimately facilitates the development of physiologically relevant engineered tissues (23–25).

To recreate tissue models *in vitro*, MECs have traditionally been isolated from mammoplasties, however, human milk has emerged as a valuable source of these cells. Recent advances in omics technologies have unveiled the presence of mammary epithelial stem and progenitor populations in breast milk. Milk MECs can be non-invasively isolated and retain key characteristics of the mammary epithelium, including the ability to form acinar structures *in vitro* (26, 27). Despite these promising prerequisites, milk MECs have to date not been applied in tissue engineering.

In this study, we present a VP lactation model that leverages cutting-edge biomaterials and 3D printing technologies and highlights the suitability of milk MECs for mammary tissue engineering. Our model paves the way for new research approaches into human lactation and provides an *in vitro* platform for exploring the impact of ECM components, as well as the effects of nutrition, hormones, and drugs on milk secretion mechanisms.

## Results

### Milk-derived MECs can be expanded and maintain lactation potential in vitro

Cells derived from human breastmilk were expanded to investigate the potential of milk MECs for *in vitro* lactation models. Human milk contained colony-generating cells with various morphologies e.g. cobble stone, refractive edges, or stratified in line with reports on epithelial cells in 2D culture (Fig. 2*A* and Fig. S1)(28, 29). Cells from all donors were expandable over at least three passages (Fig. 2*B*, Figs. S1-S2 and Table S1).

**Figure 2:**
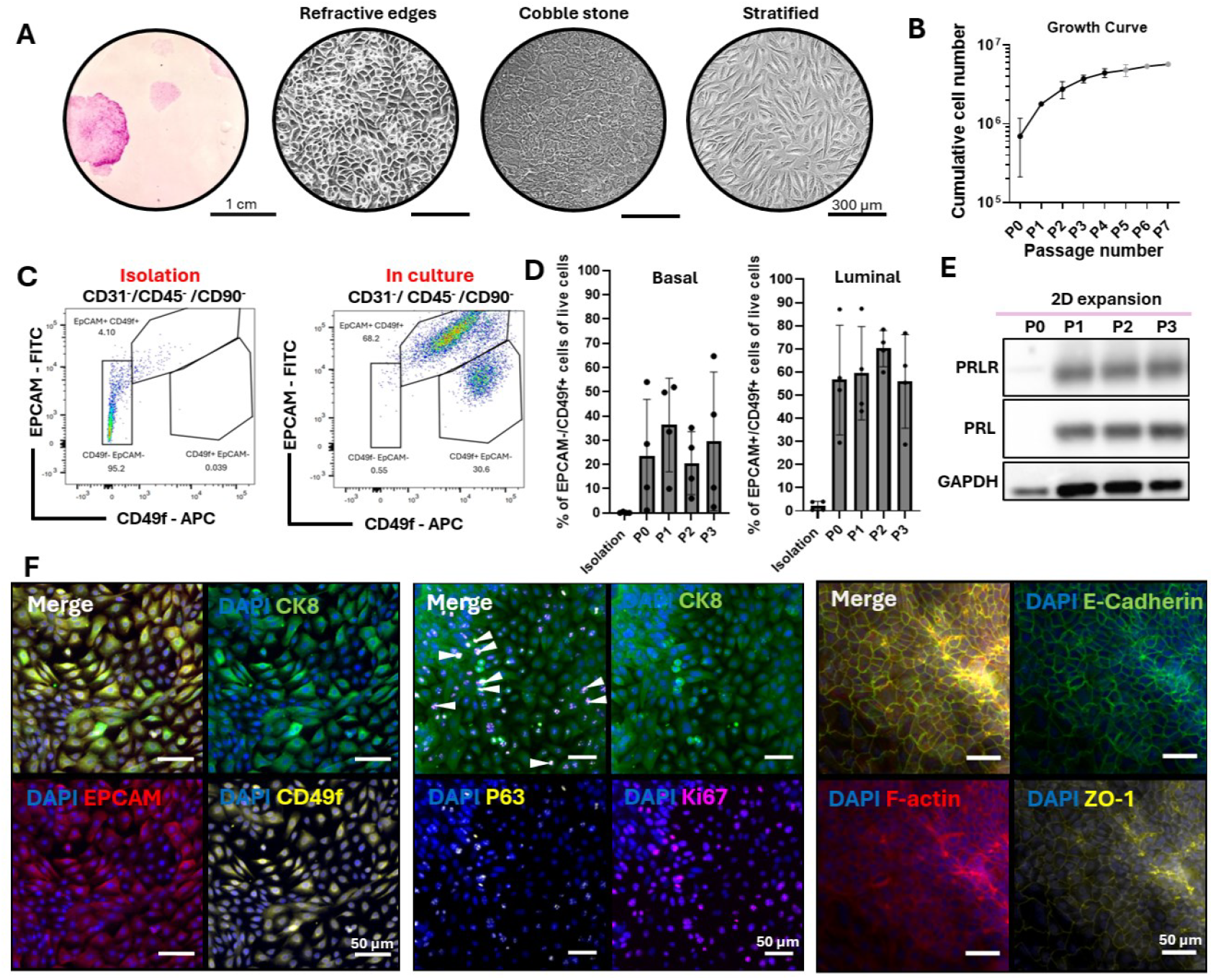
Phenotypic characterization of human milk-derived primary mammary epithelial cells (MECs) in 2D in vitro culture. A) Milk MECs post isolation (P0) stained with rhodamine B after 12 days of culture (26). Milk MECs form colonies of different sizes and cell shapes, as shown with bright field imaging. B) Milk MECs can be expanded over serial weekly passaging (x-axis, number of passages; y-axis, cumulative cell number); mean ± SD (n = 4 biological replicates represented in black, n=2 or 1 biological replicate represented in grey). C) Representative FACS plot showing the cells post isolation (epithelial cell adhesion marker negative (EPCAM-) integrin alpha-6 (CD49f-)) and during in vitro culture, demonstrating a shift towards EPCAM-/CD49+ and EPCAM+/CD49f+ epithelial cell populations. D) EPCAM-/CD49f+ basal MEC population and EPCAM+/CF49f+ luminal MEC population from 4 different donors during isolation and over 3 passages in 2D culture E) Western blot showing the presence of prolactin (PRL) and its receptors (PRLR) in the milk MEC over three passages. F) Immunofluorescent staining of milk MEC in 2D in vitro culture on Matrigel-coated polystyrene cell culture flasks with epithelial markers (EPCAM, CD49f), luminal marker (cytokeratin 8 - CK8), markers of proliferation (Ki67) and stemness (P63) with arrowhead indicating colocalization, as well as markers for tight junction formation (E-Cadherin, Zonulin-1 (ZO-1) and F-actin) (scale bar = 50 μm).

To further characterize the cells post-isolation and during expansion, flow cytometric analysis was performed and EPCAM and CD49f were used as epithelial markers to identify MECs (Fig. 2*C* and Figs. S4-S7). Freshly isolated cells predominantly lacked expression of EPCAM and/or CD49f (<10% positive for epithelial markers EPCAM and CD49f). However, following culture establishment and the first passage, the proportion of EPCAM^+^ and/or CD49f^+^ cells increased to >90%, while CD90 (mesenchymal marker) remained below 2% during passaging (Fig. 2*C* and Fig. S7). Luminal and basal MECs are crucial for milk synthesis and secretion respectively. Between the donors, the populations of basal (CD49f^+^/EPCAM^-^ and CK14^+^) and luminal MEC (CD49f^+^/EPCAM^+^ and CK8^+^) varied significantly (∼ 2% to ∼ 60% basal MECs and ∼ 30% to ∼ 90% luminal MECs), with a trend towards a higher proportion of luminal rather than basal cells (Fig. 2*D*). Taken together, these results demonstrate the presence of basal and luminal MECs during 2D culture with the preservation of epithelial phenotype over passaging (Fig. *2C-D* and Figs. S4-S7). The prolactin receptor (PRLR) is crucial for hormonal regulation and activation of downstream pathways (e.g. STAT5) that mediate lactation. Western Blot analysis of cultured milk MECs showed the retention of PRLR over several passages, which indicated the suitability for *in vitro* prolactin stimulation (Fig. 2*E*). Interestingly, milk-derived MECs expressed prolactin in a prolactin-free culture medium. This could be explained by the cells’ ability to produce prolactin through autocrine signaling (Fig. 2*E*). Presence FACS markers (EPCAM and CD49f) was reconfirmed by staining in 2D and to further determine if cultured cells were in active cell cycle, MECs were stained with Ki67 for proliferation (Fig. 2*F*). Some proliferative cells also co-expressed stem cell marker P63, which indicated the presence of basal progenitor cells (Fig. 2*F*). Additionally, milk MECs were able to form tight junctions demonstrated by Zonulin-1 (ZO-1), E-cadherin, and F-actin colocalization, a key requirement for secretory epithelial tissues to execute their function, which further indicates their potential for mammary tissue engineering (Fig. 2*F*).

### Protein-rich biomaterials can be extracted from mammary tissue

To establish a bioactive base material for bioprinted mammary scaffolds, tissue was harvested from bovine udders and decellularized (Fig. S8*A*). Hughes et al.’s work and our histological staining’s demonstrated the resemblance of the bovine to the human mammary gland with a collagen-rich stroma, in contrast with the adipocyte-rich stroma found in mice (Fig. S8*B*) (30). Additionally, fresh bovine tissue was available in large quantities, crucial given the low yield of decellularized tissue dry mass of ∼ 4% (Fig. S8*C*). After decellularization, fat, nuclei and double stranded-DNA were successfully removed (Fig. S8*D-F*). The dECM_mam_’s self-gelling properties were confirmed by a 90° thermal gelation test pre- and post-centrifugation, and gelation kinetics were assessed by thermal rheology (Fig. 3*A* and Fig. S8*F*). All three independent dECM_mam_ batches formed comparable sigmoidal gelation curves (4°C to 37 °C) and reached a plateau in about 28 min in line with previously reported studies (Fig. S8*F*)(25).

**Figure 3:**
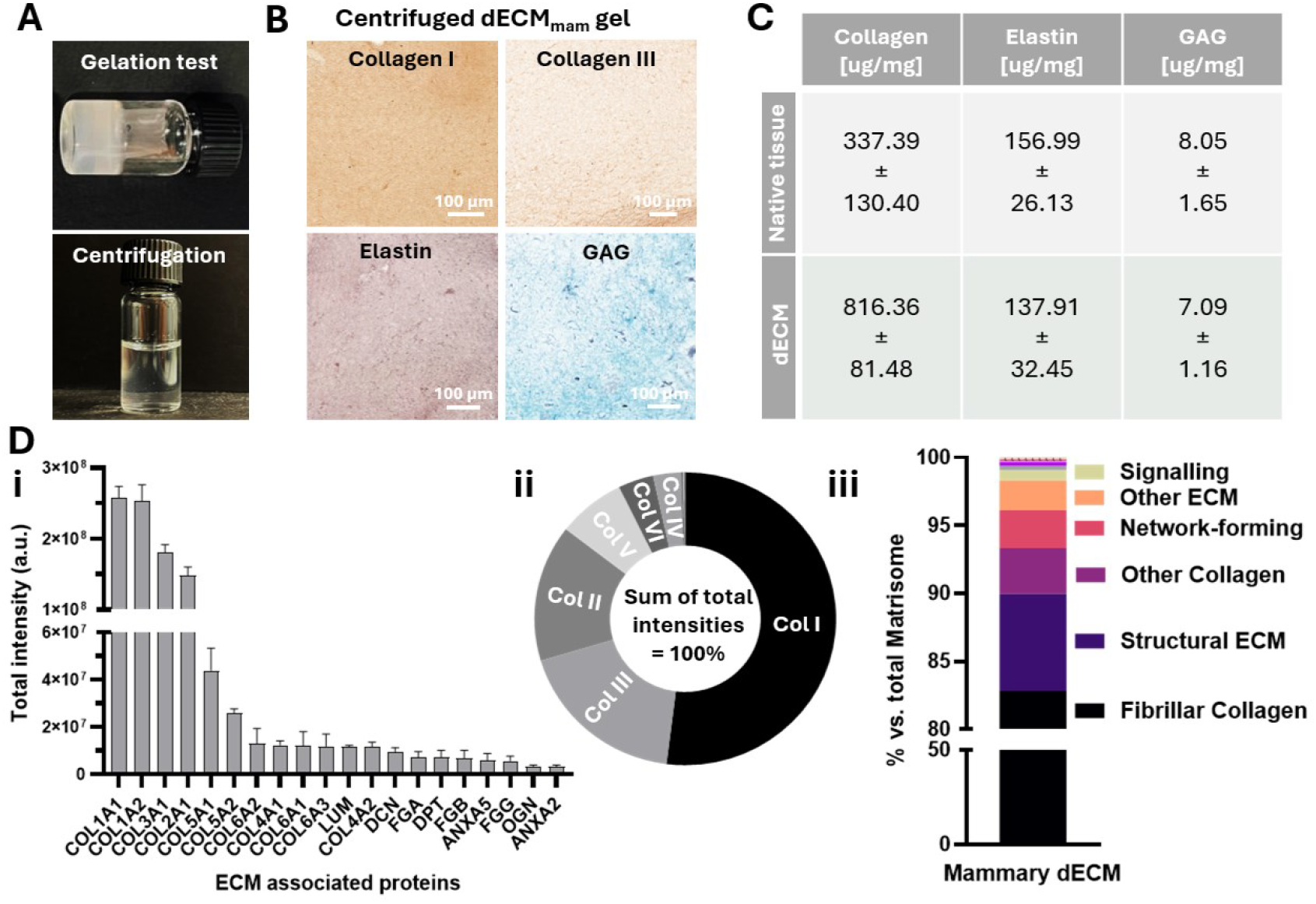
Decellularized extracellular matrix (dECM) from mammary tissue for light-based printing applications. A) The final mammary dECM (dECM_mam_) preserves self-gelling capacities (90° gelation test) post neutralization and after centrifugation (8000 rpm; 7 min) to obtain a transparent dECM_mam_ solution for light-based printing. B) Histological sections of (centrifuged) dECM_mam_ gels stained for Collagens I & III, elastin (Verhoeff’s stain) and glycosaminoglycans (GAG - Safranin-O), Scale bar 100 μm. C) Quantification of ECM proteins: collagen, elastin and GAG. Mean ± S.D. (n = 3 batches). D) Proteomic analysis of the ECM batches post decellularization employing intensity-based label-free quantification (n = 3 biological replicates): i. Top 20 matrisome proteins by total signal intensity. ii) Relative abundance of identified collagen subtypes displayed as a fraction of total collagen signal, with iii) Classification according to matrisome database (31). Fibrillar collagens were the most abundant in the decellularized ECM.

For VP the resin should be transparent to allow for light penetration and reduced scattering, hence the dECM_mam_ was further processed by centrifugation (20) (Fig. S9*A-B*). This step removed undigested particles while preserving both high and low molecular weight proteins as confirmed by SDS-Page (Fig. S2*C*). Our dECM_mam_ extraction protocol for light-based printing applications preserved key ECM components such as collagens, elastin and glycosaminoglycans (GAG) confirmed by the comparison of the native tissue to the decellularized tissue and ECM gels following trends reported in the literature (Fig. 3*B-C* and Fig. S10) (25).

For a better understanding of the native breast ECM, the residual protein content of the dECM_mam_ was determined. Mass spectrometry-based proteomic analysis of the dECM_mam_ powders pre-pepsin digestion identified 1321 proteins, of which 131 were classified as ECM or ECM-associated (Fig. 3 *Di* and Fig. S11). Collagens are essential structural and functional ECM components found in the mammary tissue and, in line with previous studies, Col I (52% of total collagen signal), III (18%), II (15%), V (7%), VI (4%) and IV (3%) were the most abundantly identified collagens in the dECM_mam_ by total signal intensity (Fig. 3*Dii* and Fig. S12*A*)(32). Identified proteins were grouped according to functional classifications derived from the matrisome database (31). Fibrillar collagens (i.e. collagens I, II, III) were the most predominant ECM components, crucial for maintaining 3D tissue structure (Fig 3*Diii* and Fig. S12*B*). To crosscheck the dECM_mam_ with other human tissues relevant for tissue engineering, a hierarchical clustering analysis of the 131 matrisome proteins was performed and compared to a draft map of the human proteome (33). The dECM_mam_ showed similarities especially with ectoderm derived tissues (i.e., epithelial and nerve tissues) (Fig. S13)

### Volumetric printed dECM scaffolds mimic human breast architecture and biophysical properties

To transform the dECM_mam_ into a light printable material the ruthenium/sodium persulfate (Ru/SPS) photoinitiator was added. Ru/SPS allowed the crosslinking of tyrosine residue-carrying proteins found in dECM_mam_ by stable di-tyrosine crosslink formation upon light exposure, without additional chemical modification (Fig. 4*A*).

**Figure 4:**
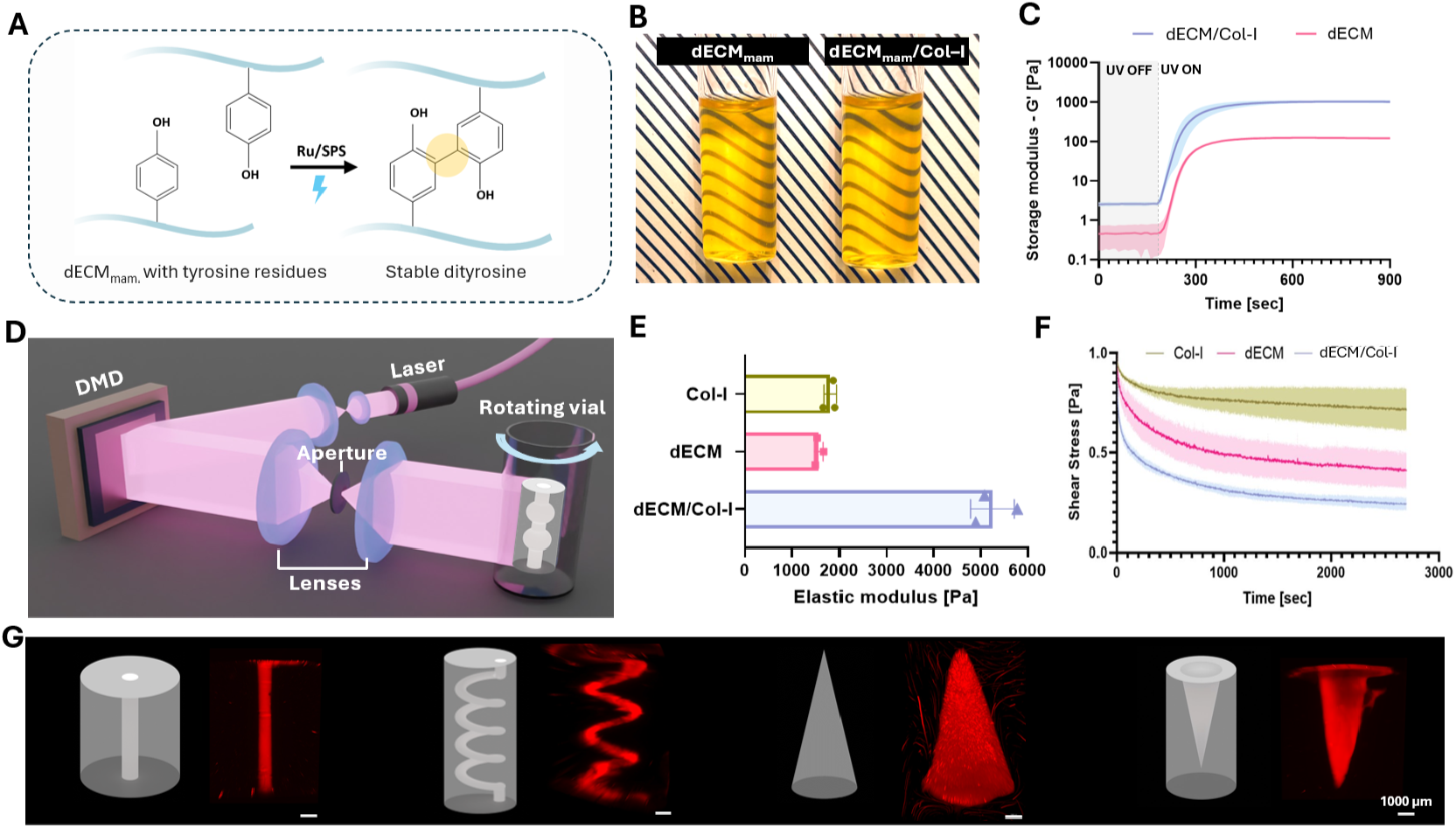
High-resolution volumetric printing (VP) of mammary decellularized extracellular matrix (dECM_mam_) for breast tissue engineering applications. A) The ruthenium/sodium persulfate (Ru/SPS) photoinitiator system allows for the formation of di-tyrosine bonds upon exposure to 405 nm laser light. B) Printing vial filled with cooled dECM_mam_ photoresin mixed with Col-I and Ru/SPS ready for light based bioprinting. C) Photorheology of the dECM_mam_ (40mg/ml) and dECM_mam_ /Col-I (40 mg/ml + 4 mg/ml) resins demonstrating a rapid gelation response upon exposure to 405 nm light. D) Schematic of VP, which allows for the creation of complex 3D shapes. E) Elastic moduli of printed and fully postcured constructs with Col-I, dECM_mam_ and dECM_mam_ / Col-I (Mean ± S.D, n = 3 technical replicates). F) Printed dECM_mam_ / Col-I constructs show slower stress relaxation responses more closely reflecting the viscoelastic behavior of the native mammary tissue (34). G) STL files depicted in grey and printed perfusable shapes (spiral and channel perfused with rhodamine acrylate (red) labeled gelatin methacrylate) and resolution tests (cone and inverted cone) printed with rhodamine-stained dECM_mam_ /Col-I resin (scale bar = 1000 μm).

Previous studies suggested that printing fidelity is strongly dependent on material and Ru/SPS concentration (22, 35). To investigate the photoinitiator’s effects on the material, Ru/SPS concentrations and ratios from 1:1 to 1:20 were iteratively tested by photorheology using a 405 nm light source (Fig. S14). Interestingly, increased photoinitiator concentrations did not correlate with faster crosslinking speeds or a higher storage modulus. This may be attributed to a limited availability of reactive tyrosine groups, whereby increased photoinitiator concentrations would not enhance crosslinking. Furthermore, higher photoinitiator levels could increase photoabsorption, that hinders light penetration and crosslinking efficiency. The gelation was further tunable by the ratio of the two photoinitiator components. A 1:10 ratio of 0.5 mM Ru to 5 mM SPS demonstrated the optimal balance of stiffness, crosslinking speed, and minimal photoinitiator concentrations to avoid cytotoxicity on cells (Fig. S14*A*).

To evaluate the effects of material concentrations, the Ru/SPS ratio was fixed at 0.5/5 mM and dECM_mam_ varied from 20 to 60 mg/ml. Beyond 40 mg/ml, additional material did not enhance the storage modulus (max.∼ 120 Pa) but slowed down the crosslinking response (Fig. S14*B*). This could be explained by particle precipitation. For mechanical property tuning of the dECM_mam_ at 40 mg/ml, collagen type 1 (Col-I) was added at concentrations between 1 and 5 mg/ml. The addition of 4 mg/ml Col-I produced a plateau storage modulus of approximately 1000 Pa, but further increases to 5 mg/ml reduced the modulus back to 450 Pa (Fig. S14*C-D*). The final photoresin optimized for mammary scaffold bioprinting consisted of 40 mg/ml dECM_mam_ and 4 mg/ml Col-I combined with 0.5 mM Ru and 5 mM SPS, with a refractive index of 1.345 (Fig. 4*B* and Fig. S15*A*). This formulation allowed to increase viscoelasticity and stiffnesses (Fig. 4*E-F* and Fig. S15*B*).

After resin optimization, dECM_mam_/Col-I spiral, cone, and inverted cone models were volumetrically printed using a light dose of 1250-1350 mW/cm^2^ for 120 seconds, achieving a positive resolution of 300 µm (Fig. 4*G* and Figs. S16-S17). Unlike constructs made from Col-I or dECM_mam_ alone, which lost their integrity within a month (Fig. S15*B-F*), the dECM_mam_/Col-I constructs maintained their shape and mechanical properties for over one month. These stable and high-fidelity structures offer a solid foundation for further exploration in mammary tissue modeling.

### Volumetrically printed dECM_mam_/Col-I alveoli support milk protein production *ex vivo*

To test biocompatibility and cell adhesion, we printed cylinders on which our milk MECs were seeded. The biocompatibility of the constructs was confirmed by high cell viability (>90%) and the ability of the MECs to completely cover the printed surface within seven days (Fig. S18*A-B*). Lactate dehydrogenase assays over a 72h period confirmed that the scaffolds were non-cytotoxic (Fig. S18*C*). The stiffness of seeded cylinders was constant over one week, demonstrating the stability and mechanical integrity of printed constructs also with cells (Fig. S18*D*).

For lactation experiments, perfusable ductal alveolar (D/A) units were designed with a channel diameter of ∼ 800 μm and alveoli diameter of ∼ 1500 μm. To test perfusion the D/A units were injected with rhodamine-labeled Gelatine methacrylate (Fig. 5*E*). Next, milk MEC were post-seeded by injection into the hollow D/A unit tube. The cells formed a continuous monolayer on the printed surface and expressed luminal (CK8) and basal (CK14) markers as well as tight junction maker ZO-1 in line with observations made during 2D expansion (Figs. 2 and 5*Bi*). Positive milk fat globule (MFG) and beta-casein staining showed droplet formation and milk protein production on our scaffolds (Fig. 5*Bi*).

**Figure 5:**
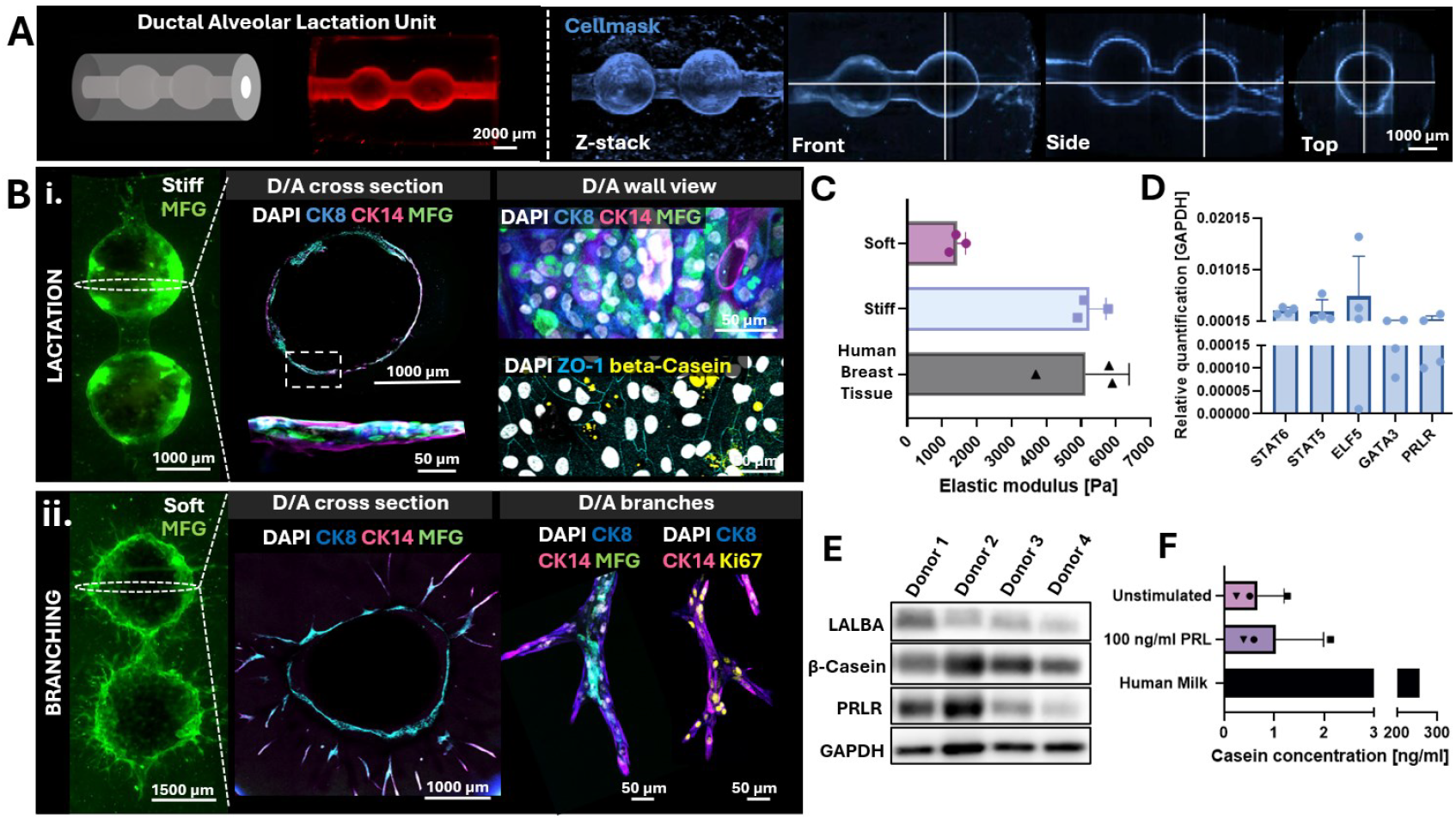
Volumetrically printed (VP) dECM_mam_/Col-I alveoli support milk MEC function ex vivo. A) Perfusable ducal alveolar units were designed using computer-aided design (grey, left). D/A units were perfused with rhodamine labeled GelMA and Cellmask labeled cells (at day 7) to show homogenous distribution of injected material and even spread of cells in the printed constructs. The white cross indicates the sections where the displayed image slices were taken from the 3D stack. B) Immunofluorescent staining of VP alveoli with milk MEC at day 7 after seeding (left: lightsheet imaging; right: confocal imaging of the same scaffold) : i) cells on stiffer constructs were stained with cytokeratin 8 (CK8), cytokeratin 14 (CK14), milk fat globule (MFG), casein and Zonulin-1 (ZO-1), to demonstrate the secretory capacities ii) cells on softer construct were stained with CK8, CK14, MFG and Ki67 to show branch elongation and proliferation. C) Elastic modulus of the softer (VP only) and stiffer (VP and post-curing) dECM_mam_/Col-I gel matched with the measurements done for healthy human breast tissue (n= 3 technical replicates for printed scaffolds, n=3 biological replicates for human tissue) D) Real time-qPCR of selected lactation relevant genes expressed by milk MEC on 3D printed constructs (mean ± SD; n = 4 biological replicates). E) Western Blot of milk-associated proteins (β-casein and lactalbumin (LALBA)), prolactin receptor (PRLR) and housekeeping gene (GAPDH) expression after prolactin stimulation of milk MEC (4 donors) seeded in the printed D/A scaffolds D) ELISA showing beta casein secretion into the media with and without the addition of prolactin and compared to human milk.

To understand the effect of material stiffness on cell behavior in our printed constructs, stiff (5000 Pa) and soft (1300 Pa) hydrogels were printed by modulating the light exposure (Fig. 5*B-C*). Cells on the softer (on the scale of Matrigel) scaffolds developed branching structures that penetrated the hydrogel with CK8 and CK14 positive MECs and MFG expression. Over a 7-day culture period, these branches grew to ∼ 750 - 1000 μm in length and continued to display proliferative potential (Ki67^+^ cells) (Fig. 5*Bii*). In the stiff gels (fabricated to match measured human breast tissue samples (3-5 kPa), also shown in Fig. 4*E*) the cells lined the printed hollow D/A units but were unable to escape the pre-defined shape. These results showcased our platform’s tunability and potential not only for lactation but also for other applications such as in mammary tissue dynamics.

For a better characterization of our lactation model in the context of signaling pathways, real time-qPCR was used to assess the expression of pregnancy- and lactation-associated genes, including Signal Transducer and Activator of Transcription 5 (STAT5), Signal Transducer and Activator of Transcription 6 (STAT6), E74 Like Transcription Factor 5 (ELF5), GATA Binding Protein 3 (GATA3), and PRLR (Fig. 5*D*). STAT5 and STAT6 transcription factors mediate prolactin and cytokines during mammary epithelial cell differentiation and lactation (36). ELF5 and GATA3 are involved in alveolar development during lactogenesis, while PRLR facilitates hormonal signaling (37, 38). Milk MECs seeded onto mammary D/A units expressed key lactation-associated genes.

The subsequent Western blot analysis of PRLR expression was in line with the RT-qPCR results in showing biological differences in PRLR expression. Beta-casein and lactalbumin (LALBA) bands reconfirmed the presence of milk proteins (Fig. 5*E* *and* Fig. S19). Interestingly, no correlation between PRLR expression levels and expression of studied milk proteins was observed in these experiments. To assess the secretion of milk components, we performed an enzyme-linked immunosorbent assay on the culture medium (Fig. 5*F*). Here, casein was detected with and without prolactin stimulation.

## Discussion

While mammary gland research was previously heavily focused on *in vivo* studies and organoid work, recent advances in light-based 3D printing offer exciting possibilities for *in vitro* breast tissue models. In this context, we developed a light-responsive biomimetic material based on decellularized mammary tissue and introduced volumetric printing to create lactating tissue scaffolds. Our printed D/A lactation units promoted adhesion, survival, proliferation, as well as the function of MECs. Non-traditional MEC sources such as human milk offer significant opportunities for lactation studies and isolated milk MECs were able to synthesize milk proteins such as beta-casein, lactalbumin, and milk fat globules in the printed lactation units.

Milk MECs were expanded in the lab while preserving their epithelial phenotype and prolactin receptors. Importantly, our FACS analysis confirmed the presence of both luminal and basal MECs which demonstrated that our culture contained the essential cell subpopulations to reconstitute mammary tissue *in vitro*. These findings contrasted with observations from fresh breastmilk cells, where previous single-cell RNA sequencing studies, along with our own FACS results, suggested that only luminal MECs are shed into the milk (∼ 4% luminal cells out of total cells and no detectable basal cells) (28). The emergence of basal cells during *in vitro* culture likely reflected cell plasticity. We showed that milk MECs can be expanded, however, under current conditions, the expansion of milk MECs remained limited to three passages. This restricted higher experimental throughput and larger milk production. Here, future efforts in milk MECs culture should consider 3D expansion techniques. Organoid research suggests that this approach enhances cell plasticity and adaptability, which can lead to more robust cell growth (40). Furthermore, the expansion of cells in their natural organization, with apical and basolateral domains, holds promise for effective restructuring into two distinct layers (luminal on top of basal MECs) also after seeding in 3D printed scaffolds, which is lacking in our current model (41, 42). The *in vivo*-like organization of the epithelium is crucial for the directional transport of milk components and for the response to lactation hormones (prolactin and oxytocin). Here, additionally, Wnt signaling and ErbB receptor pathway activation should be targeted in the future by the addition of R-spondin and Neuregulin-1 in the medium to enhance MECs’ self-renewal and further promote bilayer organization, respectively (44).

We established dECM_mam_ gels as a bioactive base material for mammary tissue engineering applications and successfully printed biomimetic breast tissue scaffolds. To provide optimal support to the cells from the moment they contact the scaffold, we could manually introduce basement membrane (BM) proteins rather than relying on the cells’ own secretion. Laminin and collagen IV are essential components of the BM, and laminin produced by basal MECs is key for progenitor self-renewal and basal-apical polarity (44, 45). Thus, the alveolar channel could be coated with a thin layer of collagen IV and laminin 111 and crosslinked with Ru/SPS upon light exposure.

On D/A units, all donor cells maintained their prolactin receptors which made them suitable for milk secretion *in vitro*. Building on this proof of concept, hormone concentrations can be further optimized, and oxytocin added for basal cell contraction upon polarization, to reach efficient lactation of milk MECs in engineered tissue models. To move towards a streamlined and dynamic system where hormones and nutrients can be readily modulated, and milk continuously collected, the volumetrically printed D/A units can be easily integrated with a peristaltic pump for perfusion (46). Perfusion will not only enhance automation and standardization, but the use of multiple D/A units in parallel would increase throughput when testing MECs’ behavior from several donors under different hormonal conditions. Furthermore, the pumping system would allow for not only continuous but also pulsatile flow patterns to recapitulate the fluctuating hormonal levels found *in vivo* and breastfeeding intervals.

Lactation potential amongst donor-derived cells varied in our study, where certain samples exhibited higher secretory capacity than others. Here, the sample collection process should be reconsidered. Observed differences may have arisen not only from biological variability but also from sample collection timing as milk composition fluctuates both throughout the lactation period and at different times of day (54). An increased sample size with set lactation time points or longitudinal studies across multiple pumping time points could offer more comprehensive insights into the characteristics of MECs throughout breastfeeding. The identification of “high potential” donor cells during identified optimal time frames would present a viable approach to maximize the *in vitro*-produced milk yield for further quantitative and qualitative milk analysis.

The materials and cells presented in this study could be adapted for multiple research questions and *in vitro* models. In future studies, a combination of natural materials like dECM or Col-I, with synthetic materials (e.g. polyacrylamide or polyethylene glycol) could further reduce (mechanical) variations of the hydrogels and offer more flexibility when tuning stiffness or viscoelasticity. This could allow us to model different states of breast development and the study of MECs-ECM interactions. Additionally, the printing resolution can be improved by the incorporation of a free radical inhibitor, such as 2,2,6,6-tetramethyl-1-piperidinyloxy (TEMPO), to enhance control over the polymerization process (47). This would allow for studies on the effects of smaller feature sizes on milk MECs’ function and the optimization of surface topography for increased milk production. Furthermore, our materials possess both photocrosslinkable and thermal gelation properties. This dual capacity suggests the use of dECM_mam_/Col-I as a thermally gelled coating in hollow fiber bioreactors, to achieve the upscaling of *in vitro* milk component production for infant nutrition down the line.

Our VP mammary model, along with its individual components such as milk MEC and dECM, provides a valuable addition to existing animal and organoid models for lactation research. Notably, the acellular nature of VP scaffolds also facilitates their broader dissemination within the field, where interested researchers could seed various cells, including MECs from healthy or diseased tissue. This flexibility holds great promise for advancing both basic and translational research in mammary gland biology and lactation.

## Materials and Methods

### Bovine tissue collection, decellularization and ECM extraction

Fresh bovine udders from milk cows were procured from the local slaughterhouse and transported to the lab immediately. The alveolar tissue was cut into ∼ 300 g pieces washed in a 5% Pen/step DI Water solution and then frozen at -20°C for short-term and at -80°C for long-term storage. For decellularization, previously published protocols for decellularization of other tissues were adapted (48). Briefly, the semi-frozen tissue was cut by hand into cubes of ∼1.5 mm^3^ and washed in MiliQ water for 2 h. The tissue pieces were washed with a 1% SDS solution for 2 h and then decellularized in fresh 1% SDS for 16h (overnight) at RT, adding 2% Pen/Strep. The pieces were then transferred to isopropanol for 5h, with frequent changes, before thoroughly washing with several MiliQ water changes. All steps were performed on a magnetic stir plate at 250 rpm. The decellularized tissue was then freeze-dried (72h) and cryomilled (Retsch) into a fine powder. The powder was enzymatically digested with 1 mg/ml pepsin in 0.1M HCL, for 48h on a magnetic stir plate (150 rpm) and then brought to neutral pH on ice (300 rpm). The neutralized dECM_mam_ was centrifuged at 8000 rpm for 7 min, freeze dried and stored at -20°C.

### Human tissue collection and preparation for mechanical measurements

Ethics approval for the study was obtained (Kantonale Ethikkomission, 2022-01844), and tissue samples were collected after written informed consent. The samples were transported to the lab on ice, where adipose tissue was dissected from the epithelial tissue. Mechanical measurements were performed exclusively on the epithelial tissue, with round sections being extracted using a biopsy punch.

### DNA, collagen, elastin and glycaminoglycan quantification

Decellularized tissue pieces were randomly chosen, and DNA was quantified using the PureLink Genomic DNA Mini Kit (Thermo Fisher). The final concentration of DNA in the samples was measured using a NanoDrop. Collagen content was measured using the QuickZyme Collagen assay kit (QuickZyme Biosciences), a Blyscan Sulfated Glycosaminoglycan Assay kit (Biocolor) was used for the glycosaminoglycans (GAGs) and an Elastin Assay kit (Biocolor) for elastin. The provider’s protocols were followed for these experiments.

### Sample preparation for proteomic analysis

Samples were taken from n = 3 bovine dECM batches (4 bovine udders pooled into one batch). Two milligrams of lyophilized material from each batch were individually combined with 100 mg of 3 mm glass beads in 1.5 mL Safe-Lock tubes (Eppendorf) and homogenized in 200 μL/mg of 6 M guanidine hydrochloride (Gnd-HCl), 100 mM ammonium bicarbonate (ABC) at power 8 for 1 minute (Bullet Blender, Model BBX24, Next Advance, Inc.). Each sample was vortexed (power 5) at room temperature (25°C) overnight. Homogenate was spun at 18,000 x g (4 °C) for 15 min and the supernatant was collected as the soluble ECM (sECM) fraction. Pellets were then treated with freshly prepared hydroxylamine buffer (1 M NH_2_OH−HCl, 4.5 M Gnd−HCl, 0.2 M K_2_CO_3_, pH adjusted to 9.0 with NaOH) at 200 μL/mg of the starting tissue dry weight. Samples were homogenized at power 8 for 1 minute and incubated at 45 °C with shaking (1000 rpm) for 4 h. Following incubation, the samples were spun for 15 min at 18,000 x g, and the supernatant was removed and stored as the insoluble ECM (iECM) fraction at −80°C until further proteolytic digestion with trypsin. All fractions were subsequently subjected to overnight enzymatic digestion with trypsin (1:100 enzyme:protein ratio) using a filter-aided sample preparation (FASP) approach and desalted during Evotip loading (described below) (49).

### LC-MS/MS analysis

Digested peptides were loaded onto individual Evotips following the manufacturer’s protocol and separated on an Evosep One chromatography system (Evosep, Odense, Denmark) using a Pepsep column (150 µm inner diameter, 15 cm) packed with ReproSil C18 1.9 µm, 120Å resin. Samples were analyzed using the instrument’s default “30 samples per day” LC gradient. The system was coupled to the timsTOF Pro mass spectrometer (Bruker Daltonics, Bremen, Germany) via the nano-electrospray ion source (Captive Spray, Bruker Daltonics). The mass spectrometer was operated in PASEF mode. The ramp time was set to 100 ms and 10 PASEF MS/MS scans per topN acquisition cycle were acquired. MS and MS/MS spectra were recorded from m/z 100 to 1700. The ion mobility was scanned from 0.7 to 1.50 Vs/cm2. Precursors for data-dependent acquisition were isolated within ± 1 Th and fragmented with an ion mobility-dependent collision energy, which was linearly increased from 20 to 59 eV in positive mode. Low-abundance precursor ions with an intensity above a threshold of 500 counts but below a target value of 20000 counts were repeatedly scheduled and otherwise dynamically excluded for 0.4 min.

### Global proteomic data analysis

Data was searched using MSFragger via FragPipe v21.1. Precursor tolerance was set to ±15 ppm and fragment tolerance was set to ±0.08 Da. Data was searched against UniProt restricted to Bos taurus with added common contaminant sequences (37,606 total sequences). Enzyme cleavage was set to semi-specific trypsin for all samples. Fixed modifications were set as carbamidomethyl (C). Variable modifications were set as oxidation (M), oxidation (P) (hydroxyproline), dioxidation (P), deamidation (NQ), Gln->pyroGlu (N-term Q), and acetyl (Peptide N-term). Label-free quantification was performed using IonQuant v1.10.12 with match-between-runs enabled and default parameters. Soluble and insoluble ECM fractions were searched separately and merged after database searching. Results were filtered to 1% FDR at the peptide and protein levels.

### Resin preparation

Col-I was resuspended at 10 mg/ml in phosphate- and glucose-rich buffer and left to dissolve overnight in the fridge. 80 mg/ml of the lyophilized dECM_mam_ were resuspended in 1X PBS on ice as stock solutions for resin preparation. 0.5mM RU and 5 mM SPS were added to the freshly prepared resin (4 mg/ml Col-I and 40 mg/ml dECM_mam_) in the dark on ice by thoroughly pipetting up and down using a viscous pipet.

### (Photo)rheology

All rheological experiments were performed using an Anton Paar MCR 302e rheometer with a 20 mm parallel plate geometry. Shear measurements were performed at a shear rate of 2% and a frequency of 1 Hz, with acquisition intervals of 10 seconds. To maintain sample integrity and prevent drying during testing, a wet tissue paper was added in the chamber. 76 µL of the sample was loaded onto the rheometer, with a gap distance of 0.2 mm.

For evaluation of thermal gelation and temperature-dependent rheological properties of the pure dECM, the lyophilized dECM powder was dissolved on ice on the same day in cooled 1X PBS and loaded onto the 4°C cooled stainless-steel floor of the rheometer. During measurement, the temperature increased to 37°C after 10 min.

For photorheology of the ECM-based resins a glass floor was used. The rheometer was combined with the Omnicure Series1000 lamp (Lumen Dynamics) with sequential 400–500 nm and narrow 405 nm bandpass filters (Thorlabs). dECM_mam_ based photoresins were prepared in the dark on ice as previously described. Measurements were left to proceed in the dark for 3 min before irradiating the sample with 405 nm light at (60%) 10 mW cm^−2^ intensity.

### Mechanical testing

Compression tests were performed on a TA.XT Texture Analyzer (Stable Micro System) with a 500g load cell. Samples were placed between the compression plates and a pre-load of 0.2 g was applied to ensure full contact of the samples with the plates. Samples were then compressed to 15% strain at 0.01mm/s. Loading and unloading curves were recorded and the compressive modulus was calculated by linear fitting the first 3% of the stress-strain curve.

### Volumetric printing

3D printing was carried out using the open format volumetric printer (Readily3D® SA, CH). The performance of the mammary photoresin was assessed with the built-in software feature for light dose tests to study the resin’s behavior in the printer’s settings, light source, and path. Prior to printing, dose tests were performed by projecting circles (ϕ = 1 mm) with increasing light intensity (784-2304 mJ/cm^2^) onto a 1 mm path length cuvette filled with the cooled dECM_mam_ based photoresin (Fig. S16). For printing, 1ml of resin was filled in the dark on ice in 10 mm Pyrex glass printing vials. To remove the non-crosslinked resin post-printing, the constructs were washed in cold 1X PBS by gently pipetting up and down. STL files for printing were created using Fusion 360. Printed constructs were postcured in a UV box to increase the stiffness of the constructs and match the mechanical properties to the one measured human breast tissue

### Milk collection, mammary epithelial isolation and culture

Human donors were recruited in line with Swiss Ethics guidelines and written informed consent was obtained from all participants by trained personnel (Kantonale Ethikkomission, 2022-02012). Human milk collected by a breast pump was immediately transported to the lab on ice. Cells were isolated by centrifugation using a previously published protocol by Twigger et al. (29). Briefly, cooled 1x PBS was added to the freshly collected milk (1:1 ratio) and centrifuged for 20 min, 780g at RT. The pellet was transferred to a fresh falcon (15 ml) and washed 3 times in 1x PBS with 1% Anti-anti by centrifuging at 480g, 5min at 4°C. The pellet was then resuspended in MECGM (PromoCell) with 10 uM Forskolin, 3 uM Rock-I (Y-27632) and 1% Pen/Strep. Cells were plated on with Matrigel-coated T25 polystyrene cell culture flasks. The milk MEC were left to attach for 5 days before changing the medium to MECGM with 10 uM Forskolin, 0.5 % Human Serum and 1% Pen/Strep (composition adapted from previously established protocols). For further cell passaging (P1 to P3) cells were cultured in MECGM with 10 uM Forskolin, 5 % Human Serum and 1% Pen/Strep. Cells were split at 80-90% confluency, using TryplE at 37°C, and reseeded at a density high density of 6000 cells/cm2 for expansion.

### Construct seeding and culture

2-4 M cells/ml were perfused into the printed alveoli. The construct was then rotated for 40 min with 4 rotations every 5 min and later 2 more rotations every 10 min. After initial cell adhesion the construct was then gently transferred to a well filled with culture medium to avoid the cells being washed out.

### Flow cytometry staining and analysis

Single-cell suspension was first incubated for 10 minutes at room temperature with BD Pharmingen™ Human BD Fc Block™ (1:50, BD, 564219) and then stained with antibody mix (table S2) in FACS buffer (PBS with 2% FBS, 0.5 mM EDTA) for 30 min at 4°C. Zombie NIR™ Fixable Viability Kit (1:1000, BioLegend, 423105) in PBS was used to discern live from dead cells. Cells were fixed in 4% PFA for 10 minutes and then washed and resuspended in FACS buffer before running the cells on a BD LSR Fortessa machine (BD FACSDiva8.0.1 software) and FlowJo^TM^ v10 software. Gating strategies are detailed in the Supplementary information. Antibody list can be found in table S2.

### Immunohistochemistry and confocal imaging

Constructs and cells were fixed in 4% PFA for 8 h or 20 min respectively, before immunofluorescent staining. The constructs/cells were washed 3 times in PBS, and blocked with 5% bovine serum albumin (BSA, Millipore Sigma) with 0.2% Triton-X100 in PBS for 35 h at room temperature. The constructs were then incubated with primary antibodies (table S2) in BSA-PBS overnight at 4°. On the next day, the samples were washed 3 times in PBS and incubated with the secondary antibody solution (supplementary information table 2) and DAPI diluted in BSA-PBS for 2h at RT. Before imaging samples were washed 3 times in PBS and imaged using a FVOlympus 3000 confocal microscope. Antibody list can be found in table S2. Used antibodies can be found in tables S3 and S4.

### Western Blotting

Milk MEC were grown on 3D printed constructs for 2 days and were stimulated with prolactin for another 5 days prior to protein isolation. Cells were lysed in RIPA buffer supplemented with protease inhibitors (Sigma-Aldrich P1860-1ML, Burlington, Massachusetts, USA) and then centrifuged for 10 min at 12,000 × g. Total protein contents were calculated using Pierce 660 nm Protein Assay (Thermo Scientific™, 22660, Waltham, Massachusetts, USA). 10 µg of proteins were mixed with NuPAGE™ Sample Reducing Agent (Invitrogen™ NP0004, Waltham, Massachusetts, USA) and NuPAGE™ LDS Sample Buffer (Invitrogen™ NP0007, Waltham, Massachusetts, USA) and denatured for 10 min at 80°C using a thermocycler. Samples were run on a NuPAGE™ 4–12%, Bis–Tris, 1.0–1.5 mm, Mini Protein Gel (Thermo Scientific™ NP0321BOX, Waltham, Massachusetts, USA) and transferred onto a nitrocellulose membrane. The membrane was incubated over night with primary antibodies (listed in the supplementary table 2) at 4°C. Membranes were washed with PBS supplemented with 0.1% Tween and incubated with a secondary HRP-conjugated goat anti-rabbit or anti-mouse IgG antibodies (Table S5). The HRP signal was detected with WesternBright ECL HRP substrate (Advansta K-12045-D20, San Jose, California, USA) and imaged with a FUSION FX6 EDGE Imaging System (Witec, Sursee, Switzerland).

### RNA extraction and real-time qPCR

Milk MEC were cultured in two technical duplicates for each biological replicate on 3D printed constructs for 5 days. Cells were incubated for 10 min in BLT buffer and then mixed with 70% EtOH before transfer to the RNeasy mini kit (Qiagen) column to extract the total RNA. An A260/280 ratio of between 1.8 and 2.1 was accepted as adequate quality for the RNA samples. 100 ng of RNA was retrotranscribed with GoScript Reverse Transcriptase kit (Promega A5003, Madison, Wisconsin, USA). cDNA was diluted 1:5 with RNAse-free water. RT-qPCR was performed with GoTaq qPCR Master Mix (Promega A6002, Madison, Wisconsin, USA). Reactions were run on a QuantStudio 3 96-well 0.1 ml Block Real-Time PCR System (Applied Biosystems™, Waltham, Massachusetts, USA). Samples were analyzed using amplification and melt curves. Ct values were normalized to GAPDH. All RT-qPCR primer pairs designed for this study were bought through Microsynth AG (Balgach, Switzerland) and can be found in table S6.

### Light sheet microscopy

An axially scanned light sheet microscope (MesoSPIM, V4) was used to image the volumetrically and Fyrefly printed constructs. The constructs were transferred to a 4 mm glass cuvette with miliQ water and mounted onto a custom 3D-printed sample holder and submerged in a quartz chamber filled with miliQ water, mounted onto the MesoSPIM microscope stand. For imaging, a macro-zoom system (Olympus MVX-10) and 2x air objective (Olympus MVPLAPO1x) with adjustable zoom were used. Voltage adjustments using the electrically tunable lens (ETL) were performed for each run. Step size was chosen from 5 – 50 µm.

### Quantification and statistical analysis

In all presented experiments three or more biological replicates were performed. The exact sample size (n) is indicated in the figure legends. Data analysis was performed in Excel, Matlab and GraphPad Prism software (GraphPad) 9.0. Unless otherwise specified the mean ± SD and statistical significance was determined using specified test (figure legends). P values < 0.05 are considered significant.

## Acknowledgments

We first want to thank the participating mothers and their children without whom this research would not have been possible. We thank Shannon Kelleher, Philipp Fisch, Mara Saenz de Juano Ribes as well as all the members of the lab for their inputs. We want to thank Giulia Silvestrelli and Susanne Ulbrich for helping with the procurement of the bovine tissue and for sharing reagents. We thank Thomas Biesgen for providing the human breast tissue. We greatly appreciate Jakub Janiak’s rendering of the volumetric printer. We thank Colette Bigosch for her help with the ethics approvals. The authors gratefully acknowledge ScopeM for their support and assistance in this work. Lightsheet imaging was performed with equipment maintained by the Center for Microscopy and Image Analysis, University of Zurich. We thank the Flow Cytometry Core Facility of the ETH Zurich for cell sorting and/or support with flow cytometric analysis.

## Author contributions

M.Z.W. and A.H. designed the study. A.H. carried out the experiments. K.B. performed the flow cytometry data analysis. M.M., A.S., K.H. and K.C. provided mass spectrometry data. K.C. shared decellularization protocols and helped set up the procedure for the mammary tissue. P.C shared reagents and assisted with the volumetric printer set up. A.H. and M.Z.W. wrote the manuscript. All authors helped by shaping the research, presentation of results and manuscript by providing critical feedback. M.Z.W. acquired the funding.

