## Supplementary Material for "Volumetric printed biomimetic scaffolds support *in vitro* lactation of human milk-derived mammary epithelial cells"

Amelia Hasenauer *et al.*

*Marcy Zenobi-Wong,

**This PDF file includes:**

Figures S1 to S19
Tables S1 to S6

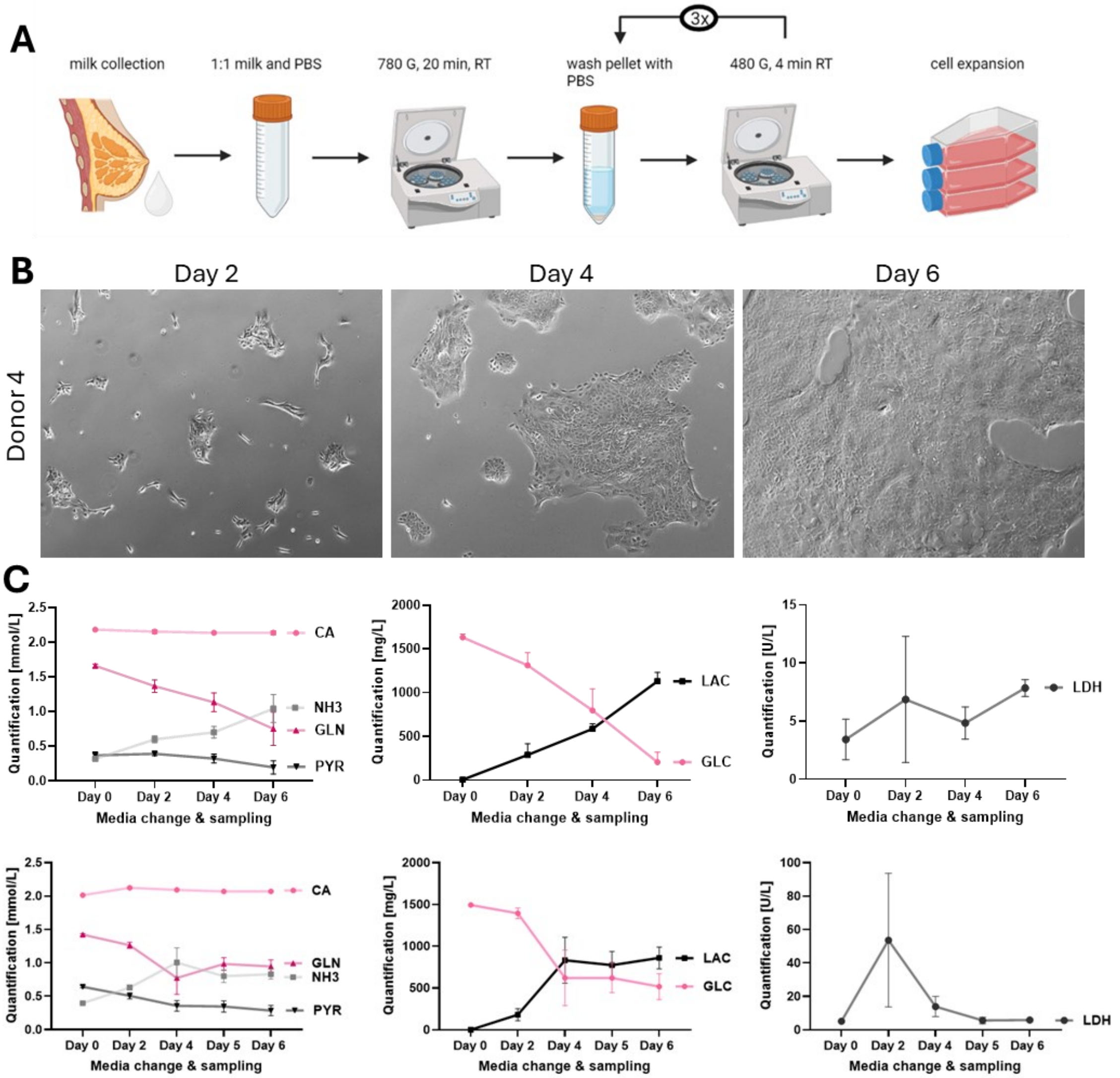


**Figure S1: Milk cell isolation and culture**

A) Human milk cells were isolated based on centrifugation. B) Brightfield images of milk isolated cells at P3 in mammary epithelial cells growth medium (MECGM, Promocell) with 5 % human serum and 3 nM forskolin in hypoxic conditions with media changes every two days. C) Analysis of the media at every media change over the culture period. Measurements of levels of calcium, ammonia, glutamine, pyruvate, lactate, glucose and lactate dehydrogenase using the Cedex media analyzer.

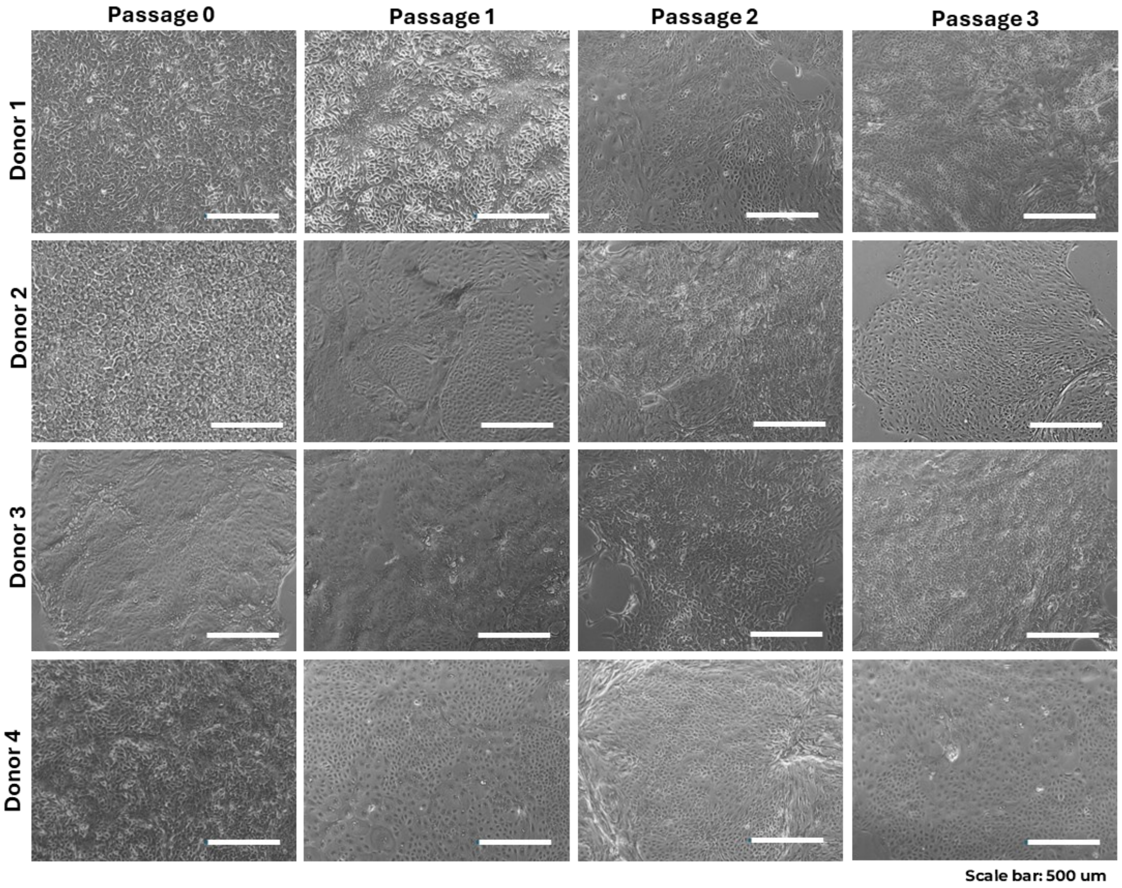


**Figure S2: Milk cell expansion in 2D over serial weekly passaging**
Expansion of milk derived cells from 4 different donors over serial weekly passaging. Brightfield images of cells at day 7 at 85-90 % confluency showing different colony types: refractive edges, cobble stone and stratified. Scale bar 500 um.

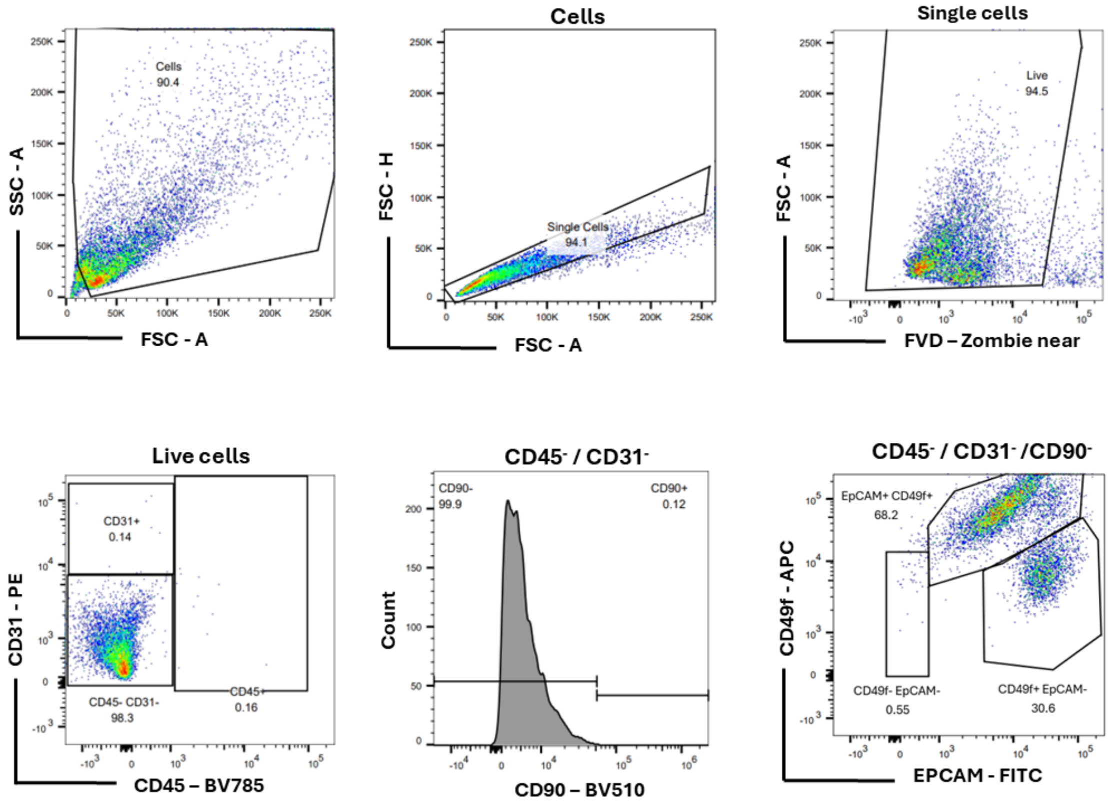


**Figure S3: Flow Cytometry gating strategy for milk derived cells**

Flow cytometric panel was designed using previously established protocols for breast tissue derived cells to identify mammary epithelial cells and separate in basal and luminal epithelial phenotypes.

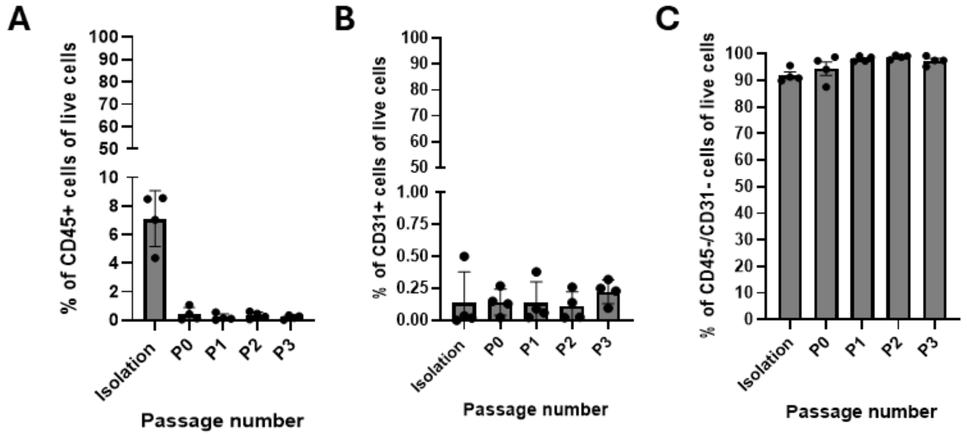


**Figure S4: Overview of the live cell populations from isolation and over 3 passages analyzed by Flow Cytometry.**

A) Percentage of CD45 expressing cells; B) percentage of CD31 expressing cells and C) percentages CD45 and CD31 negative cells during isolation and over three passages (n=4 donors; mean + SD)

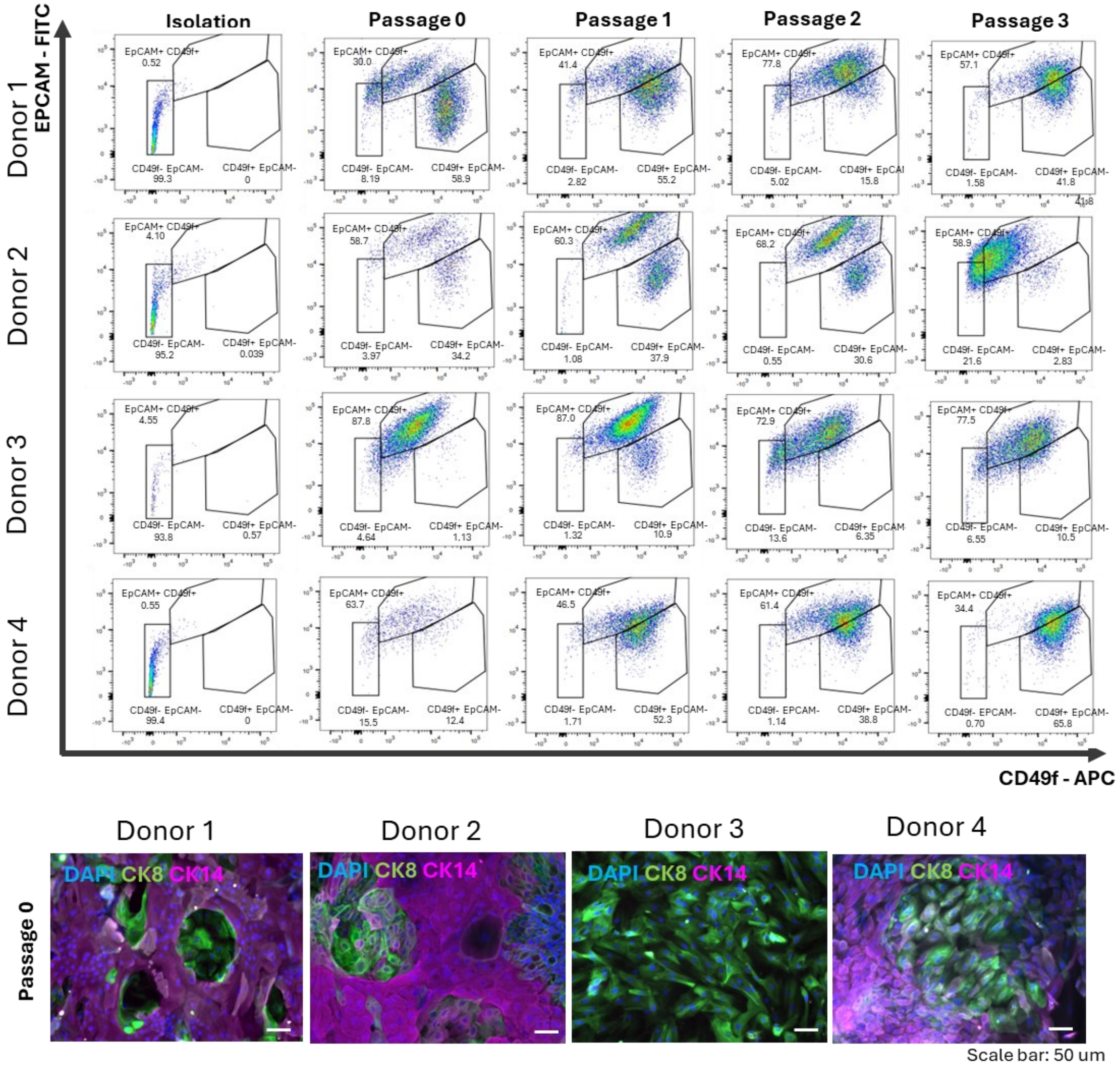


**Figure S5: Mammary epithelial populations over 3 passages analyzed by Flow cytometry**
Top: Milk derived cells were cultured in vitro and split at 85-90 % confluency. Flow cytometric analysis shows the mammary epithelial cells populations over serial weekly passaging. The cells display a luminal, basal or hybrid luminal basal phenotype. (n= 4 biological replicates)
Bottom: Immunofluorescent staining for CK8 and CK14 markers in P0 milk MEC from 4 donors.

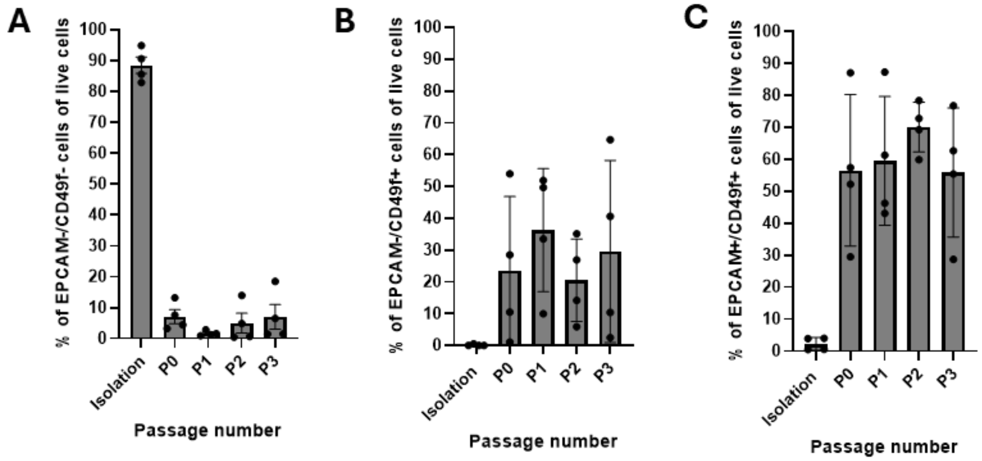


**Figure S6: Overview of the CD45⁻ / CD31⁻ /CD90⁻ cells identified by Flow Cytometry displaying epithelial phenotypes.**

A) Percentage of EPCAM and CD49f negative cells (non-epithelial phenotype); B) percentage of EPCAM negative and CD49 positive cells (basal phenotype) and C) percentages double positive cells for EPCAM and CD49f (luminal phenotype) cells during isolation and over three passages (n=4 donors; mean + SD)

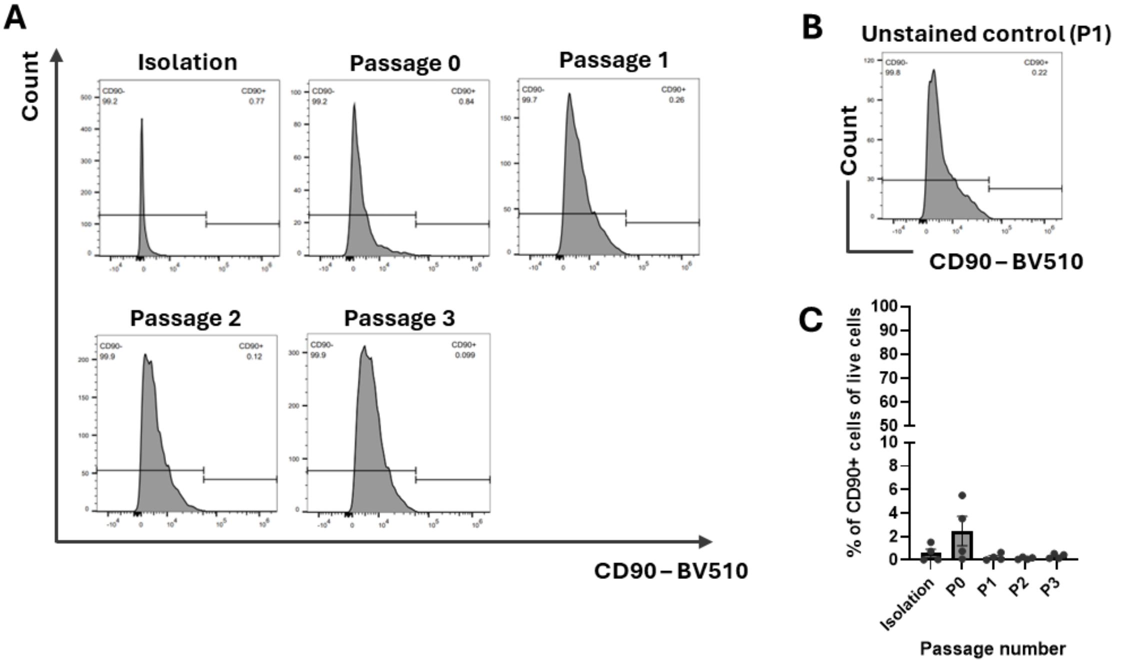


**Figure S7: Absence of CD90 expression indicating the absence of mesenchymal cells during 2D in vitro culture.**

A) CD90 expression of $CD45^-$ / $CD31^-$ cells during isolation and 2D culture after 7 days of culture split at 85-90 % confluency. Y-axis: event count; X-axis: CD90 – BV510 signal. B) Unstained control of isolated mammary epithelial cells at passage 1. Y-axis: event count; X-axis: CD90 – BV510 signal. C) CD90 expression of $CD45^-$ / $CD31^-$ cells determined by Flow cytometric during isolation and 2D culture after 7 days of culture split at 85-90 % confluency. (n=4 donors; mean + SD)

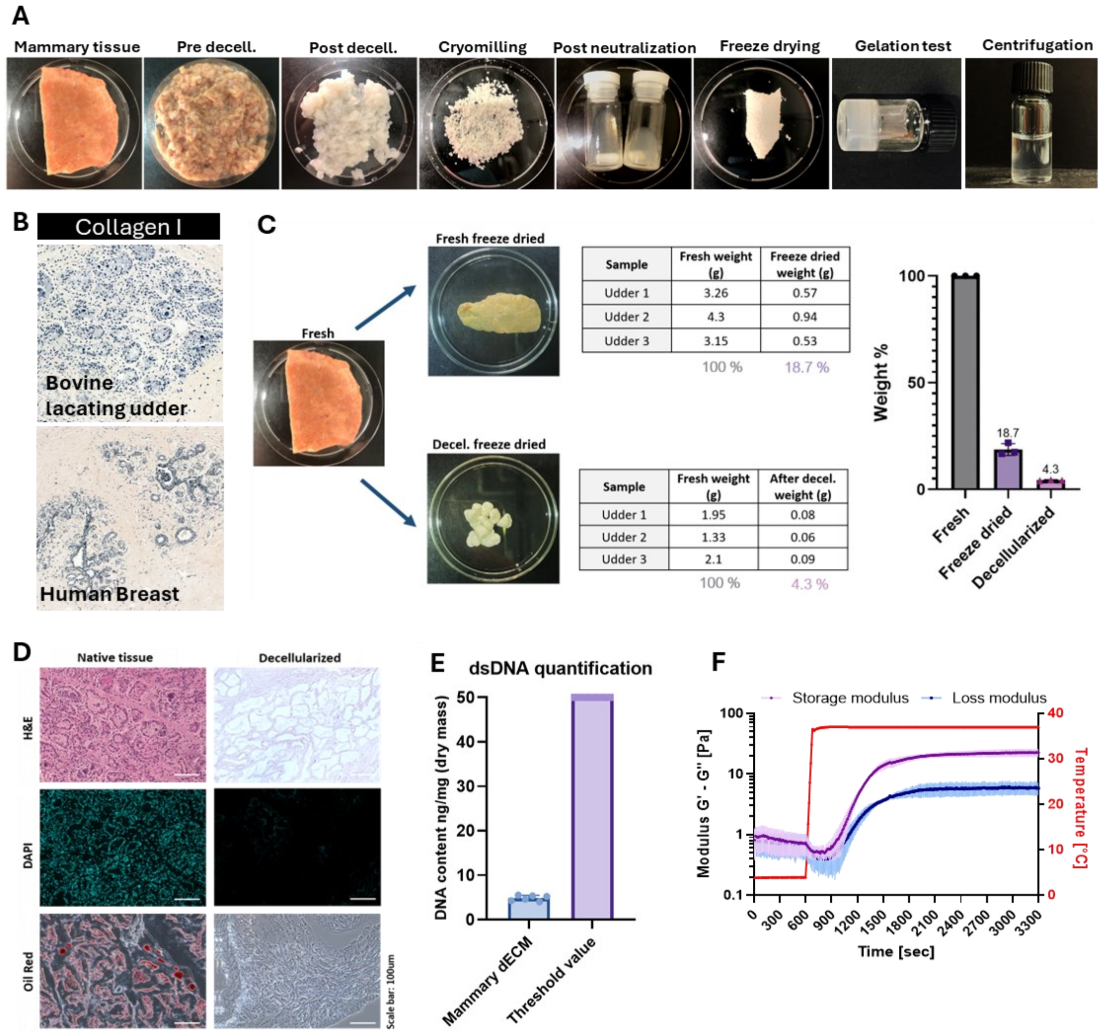


**Figure S8: Tissue decellularization**

A) Decellularization process: Bovine mammary tissue was (i) harvested, (ii) decellularized, (iii) freeze dried and cryomilled, (iv) digested and neutralized , and (v) freeze drying again for long term storage. Weight percent of freeze dried and freeze dried and decellularized tissue vs. fresh bovine mammary tissue (n= 3 biological replicates). B) Col-I stainings of histological sections of bovine and human mammary tissue. C) Weight before (fresh tissue) vs. after decellularization and freeze drying. D) Histological comparison of tissue structure (H&E), of cell (DAPI) and lipid (Oil RED) content between the native vs. decellularized bovine mammary tissue. E) Double stranded DNA quantification of the decellularized mammary tissue vs. the published threshold value (50 ng/ml) indicating a successful cell removal. F) Rheological profile of the centrifuged $dECM_{mam}$ batches resuspended at 40mg/ml in cold PBS.

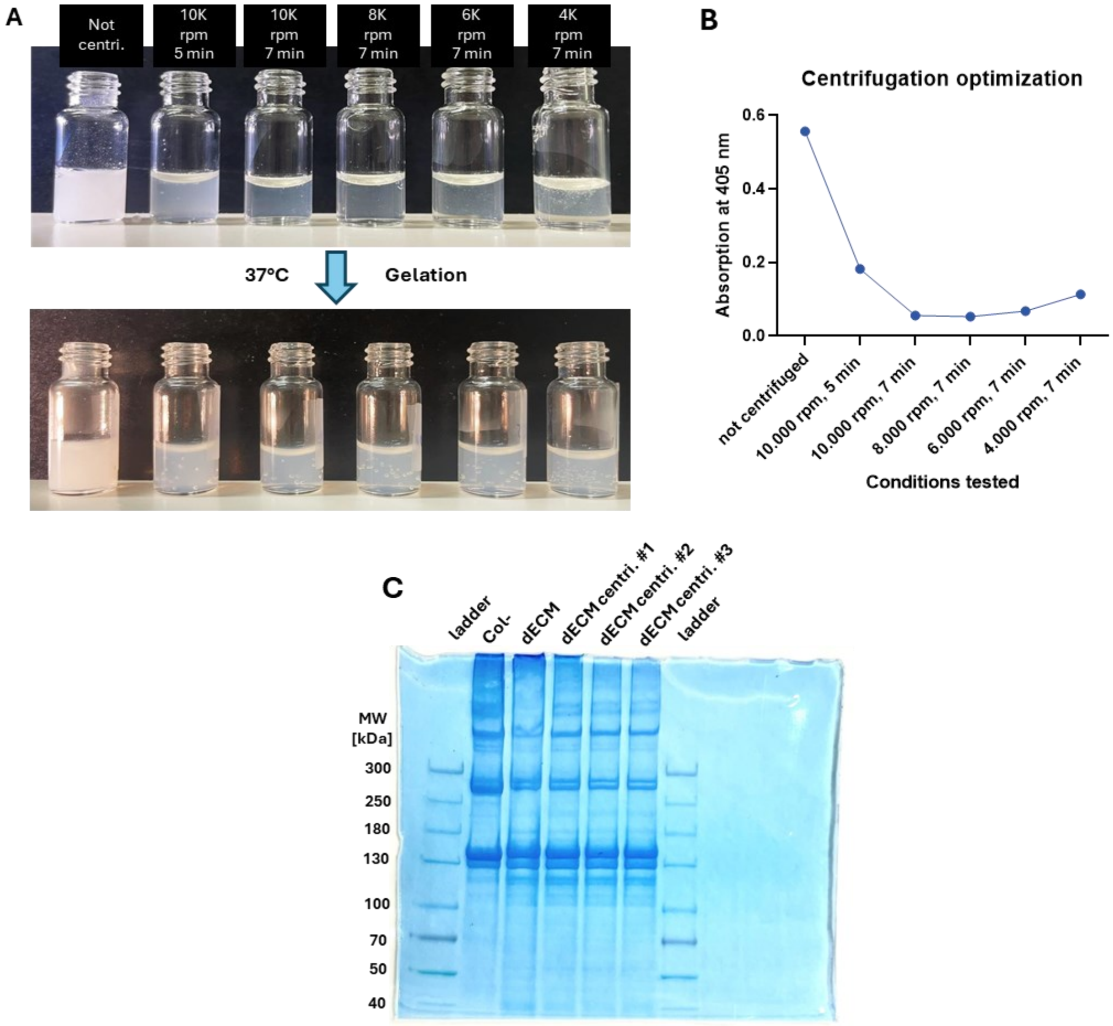


**Figure S9: Optimization of optical properties by centrifugation**

A) $dECM_{mam}$ (40mg/ml) centrifuged at 4°C at different speeds and time, maintaining its self-gelling capacities when placed at 37°C. B) Measurement of the absorption of the $dECM_{mam}$ centrifuged under different conditions at 4°C in the liquid state. C) Biochemical composition of the $dECM_{mam}$ gels via SDS Page compared to Col-I

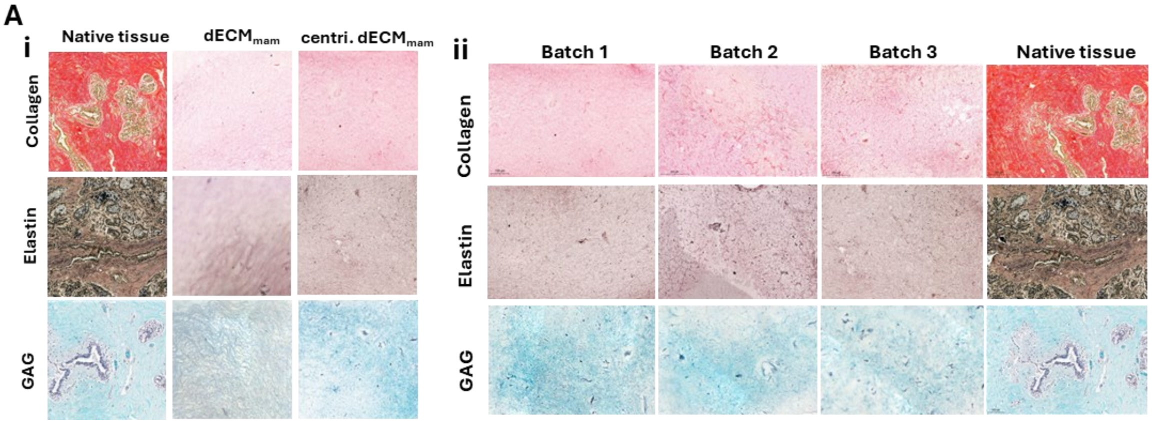


**Figure S10: Histological identification of main functional components of the ECM**

A) i. Histological sections of fixed native tissue and (centrifuged) $dECM_{mam}$ gels stained with Picrosirius Red, Verhoeff's and Safranin-O for collagen, elastin and glycosaminoglycans, respectively, showing the presence of these functional components in the native tissue, as well as in the $dECM_{mam}$ gels pre and post centrifugation resuspended at 40 mg/ml in PBS, ii) and comparison of the three centrifuged $dECM_{mam}$ batches resuspended at 50 mg/ml to the native tissue.

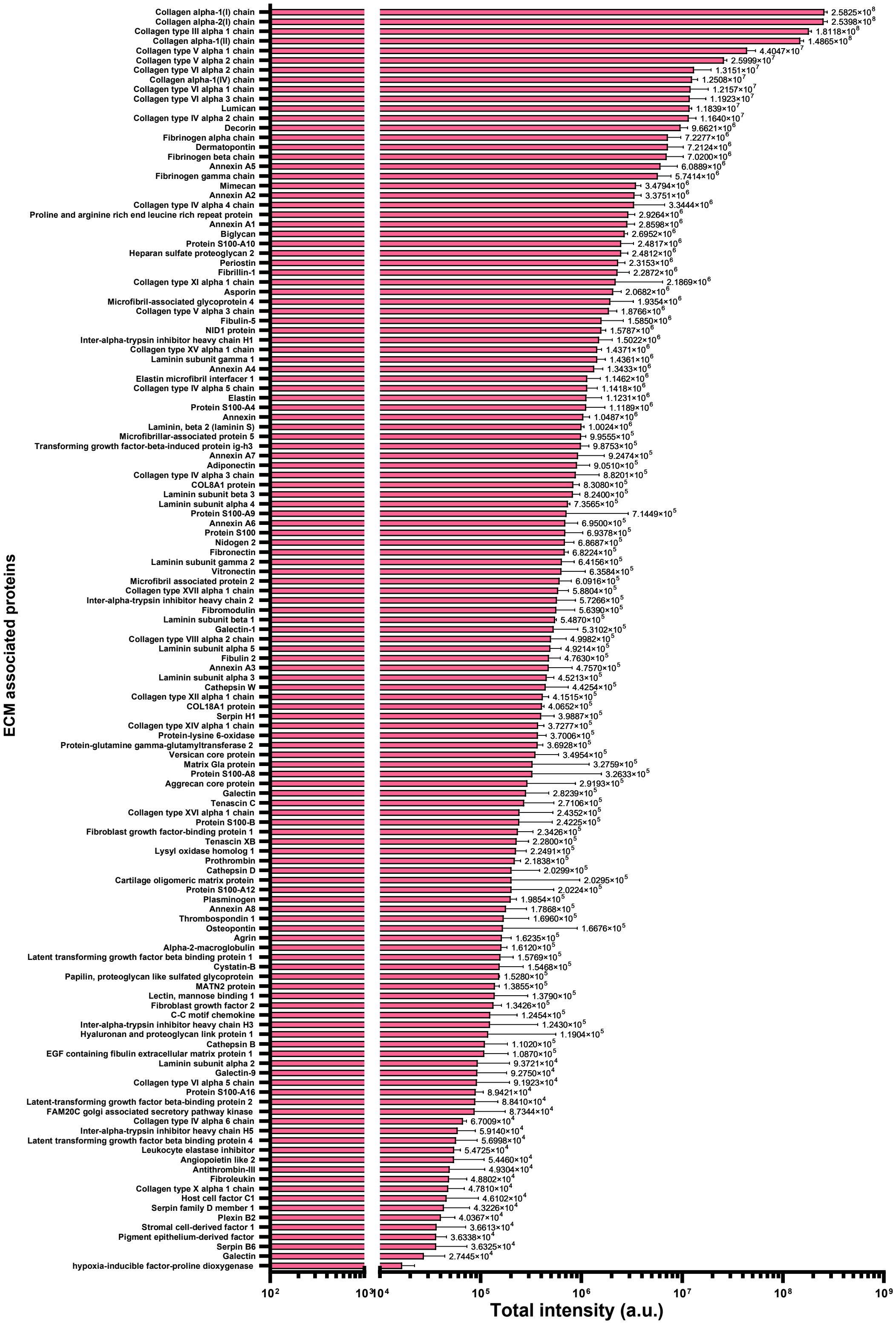

**Figure S11: Relative abundance of all 132 ECM proteins identified by mass spectrometry**

A

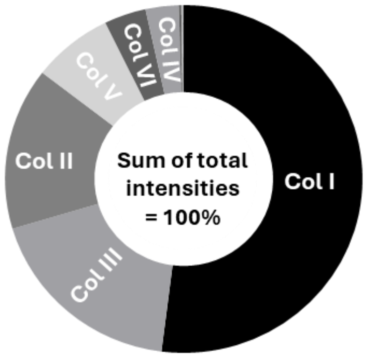


| Collagens | Distribution [%] |
|---|---|
| Col I | 51.99 |
| Col III | 18.39 |
| Col II | 15.09 |
| Col V | 7.30 |
| Col VI | 3.79 |
| Col IV | 3.00 |
| Col XI | 0.22 |
| Col XVII | 0.06 |
| Col VIII | 0.05 |
| Col XII | 0.04 |
| Col XIV | 0.04 |
| Col XVI | 0.02 |
| Total | 100.00 |

B

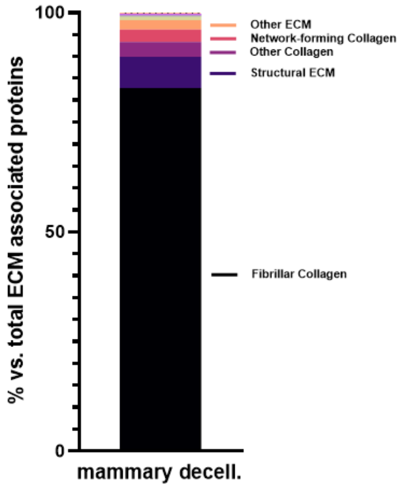


| ECM associated proteins | Distribution [%] |
|---|---|
| Fibrillar Collagen | 82.82 |
| Structural ECM | 7.13 |
| Other Collagen | 3.37 |
| Network-forming Collagen | 2.80 |
| Other ECM | 2.17 |
| Signalling | 0.79 |
| Protease Inhibitor | 0.30 |
| Matricellular | 0.27 |
| Proteases | 0.11 |
| FACIT Collagen | 0.09 |
| ECM regulators (ECM enzymes) | 0.09 |
| Membrane Collagen | 0.05 |
| ECM PTM Modifier | 0.01 |
| Total | 100 |

C

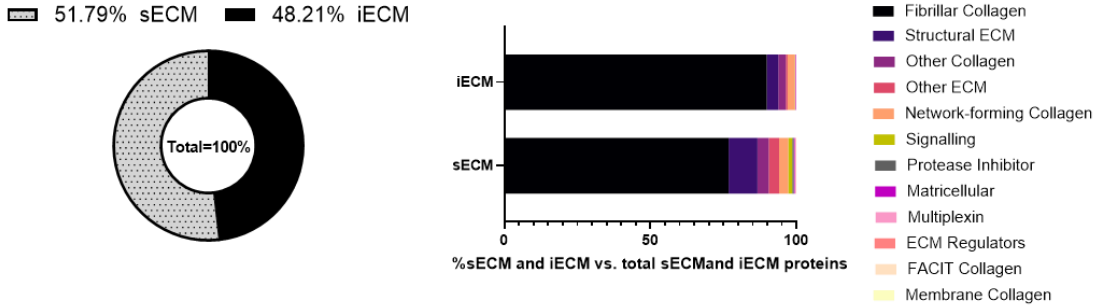


**Figure S12: Proteomic analysis**

A) Abundance of ECM-associated proteins based on DAVID gene ontology functional groups. Fibrillar collagens are the most abundant group found in the bovine mammary decellularized matrix. B) Percentage of proteins identified within the sECM, and iECM fractions of bovine decellularized mammary gland according to the DAVID gene ontology functional group classification.

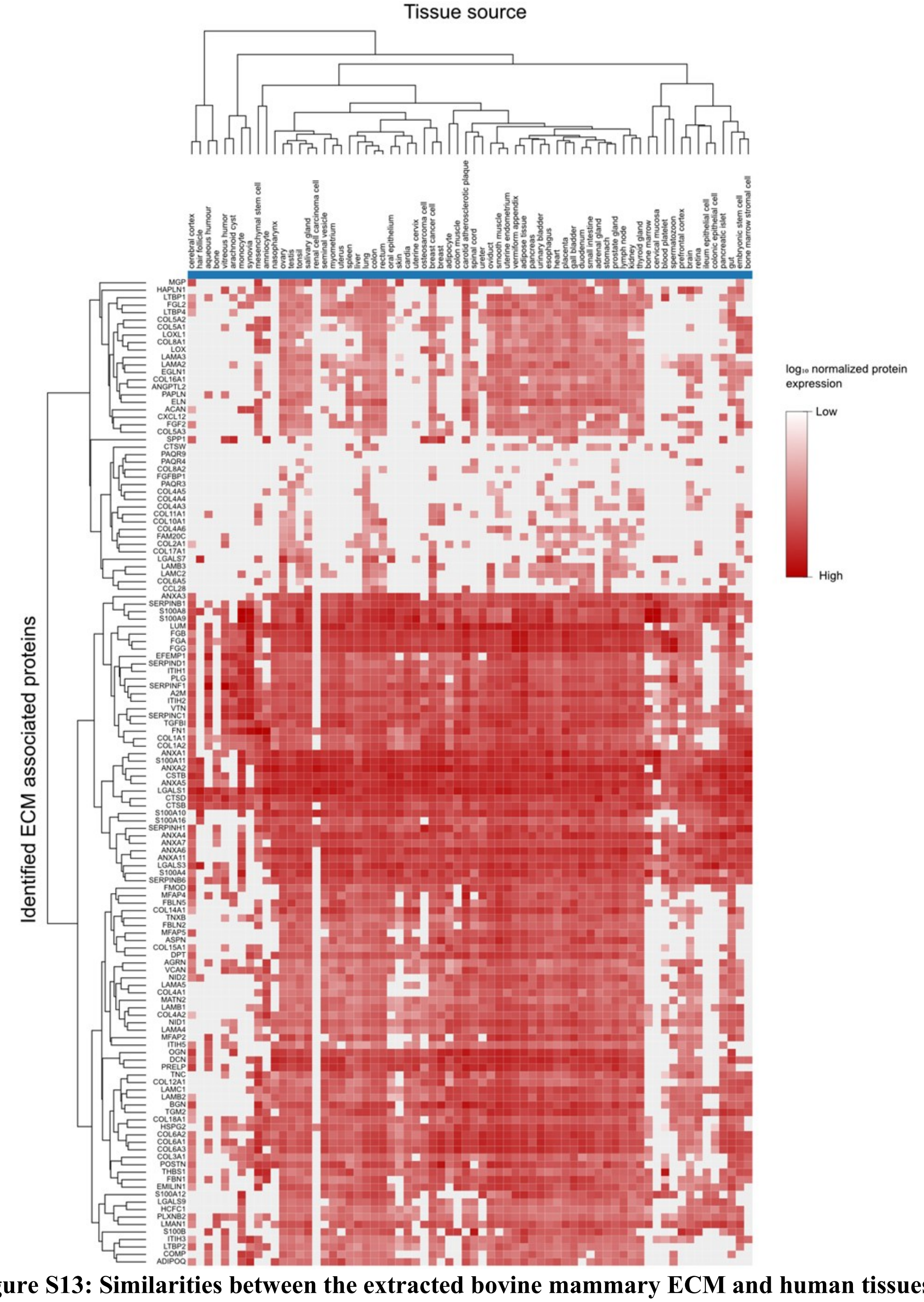


**Figure S13: Similarities between the extracted bovine mammary ECM and human tissues.** Heatmap of the hierarchical clustering analysis of the 131 ECM associated proteins identified in the decellularized bovine tissue and compared to the human tissue data from a draft map of the human proteome made publicly available proteomicsdb.org. The analysis was restricted to relevant tissues for regenerative and tissue engineering applications. The 131 identified proteins were predominantly found in ectoderm derived tissue (e.g. epithelial and nerve tissues).

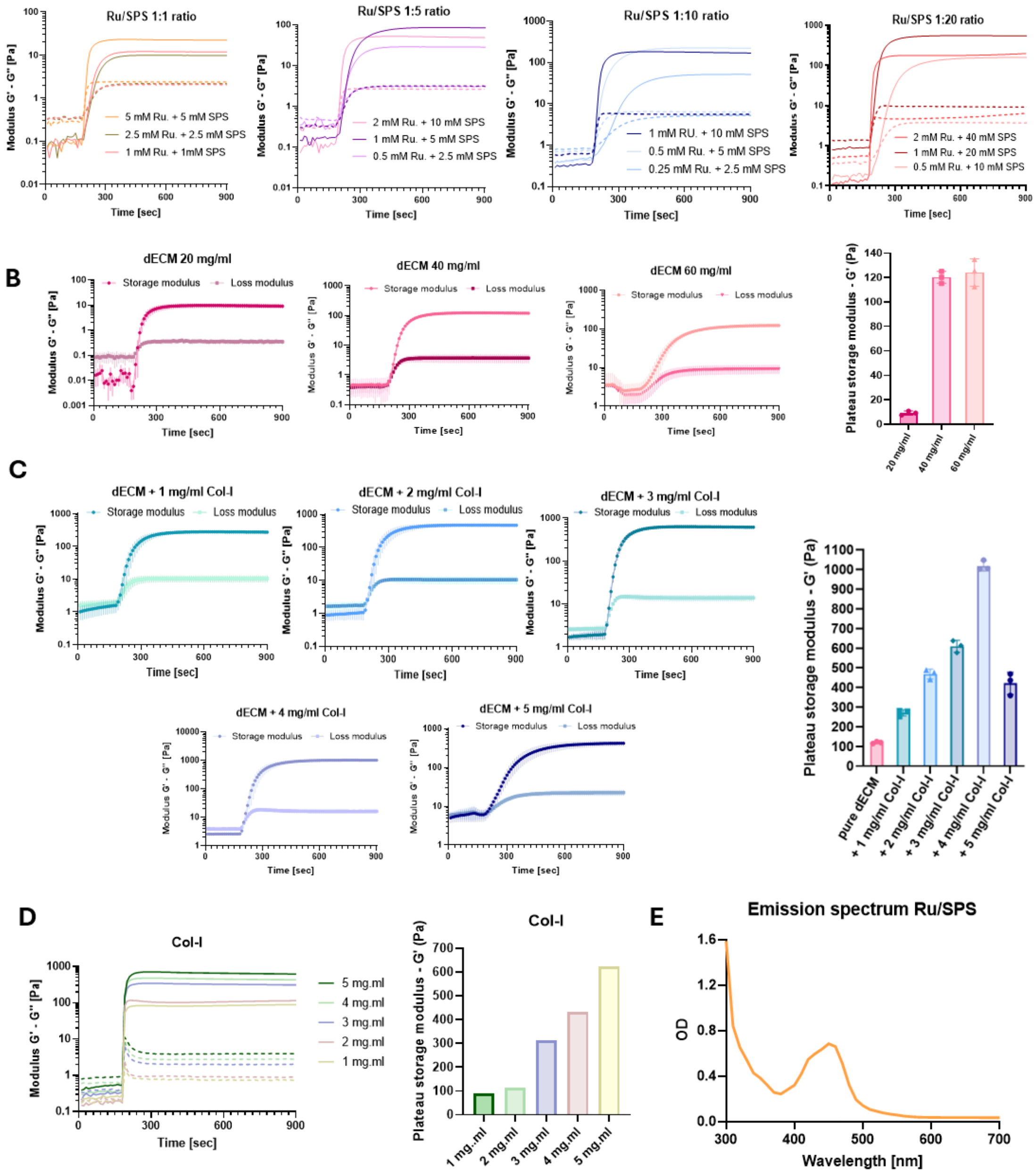


**Figure S14: Optimization of resin composition and photoinitiator ratios**

A) Photorheological profile of the $dECM_{mam}$ (40mg/ml) + Col-I (4mg/ml) resin with different Ru/SPS concentrations and ratios. B) Photorheological profile of different 3 $dECM_{mam}$ concentrations with 0.5mM RU/5mM SPS (n= 3 $dECM_{mam}$ batches) C) To make the 3 $dECM_{mam}$ resin stiffer Col-I was added to the resin in different concentrations (1 to 5 mg/ml); 0.5mM Ru/ 5mM SPS) (n= 3 $dECM_{mam}$ batches) D) Photorheological characterization of Col-I at different concentrations (1 to 5 mg/ml); 0.5mM Ru / 5mM SPS. E) Emission spectrum of the used Ru/SPS photoinitiator system with the absorption maximum at 450 nm.

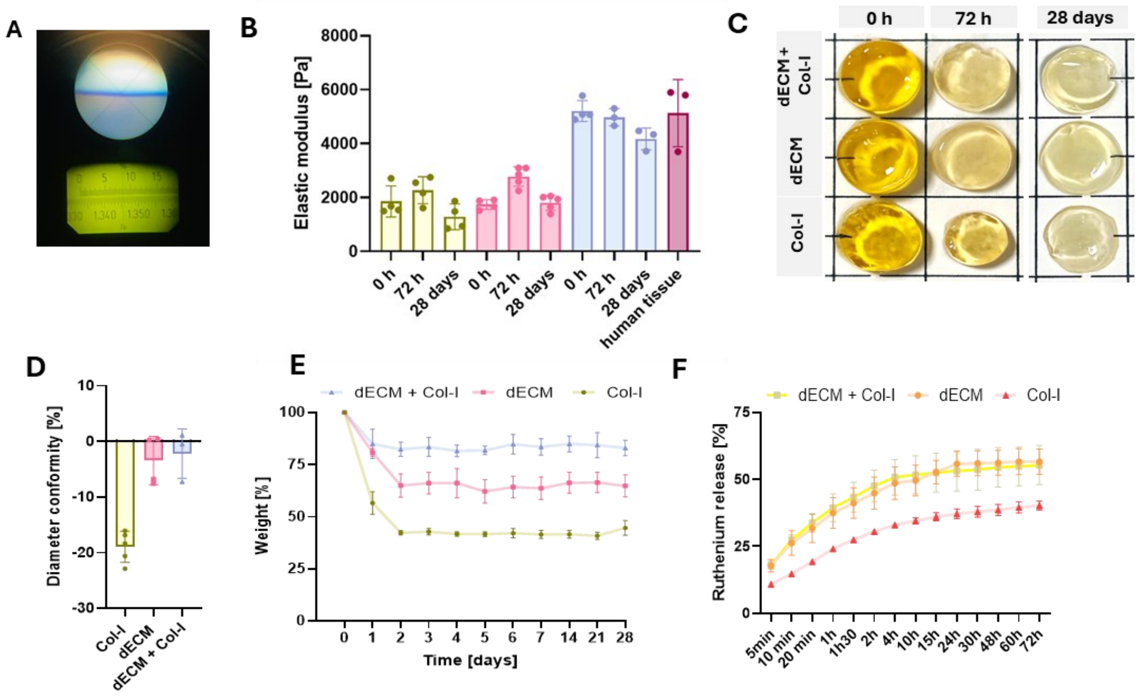


**Figure S15: dECM/Col-I photoresin for mammary tissue engineering.**

**A)** Refractive index measurement of the $dECM_{mam}$ / Col-I resin of 1.345. B) Assessment of the stability of volumetrically printed (VP) disks, over the time span of one month at RT in PBS on a shaker 150 rpm. C) The $dECM_{mam}$/Col-I was most stable as also shown by the analysis of the disk diameters. D) Elastic moduli of the Col-I (4mg/ml), $dECM_{mam}$ (40 mg/ml) and $dECM_{mam}$ + Col-I (40 mg/ml + 4 mg/ml) VP printed disks post printing, after 3 days, and after one month at RT, in PBS on the shaker 150 rpm; E) Measurement of the weight VP disks over one month F) Analysis of the RU/SPS release from the VP printed disk by photometric analysis of the PBS solution.

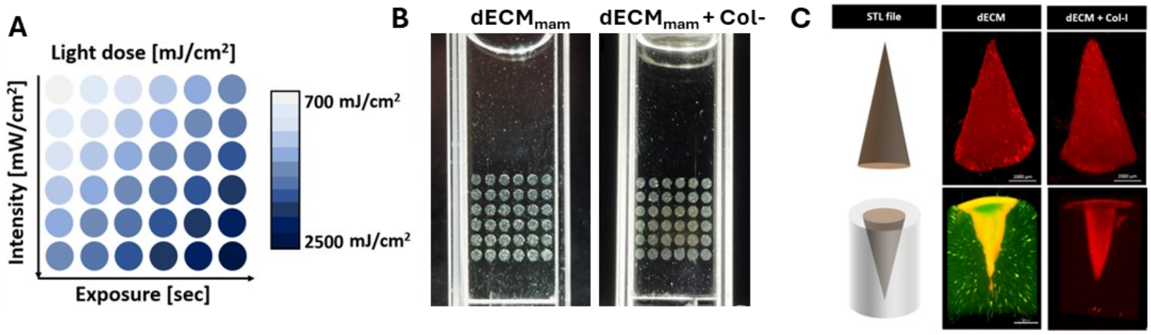


**Figure S16: Volumetric printing of dECM and dECM/Col-I**

A) Dose test set-up from 700 to 2500 mJ/cm2 to determine the printing window. B) $dECM_{mam}$ and $dECM_{mam}$/Col-I were mixed with 0.5mM RU/5mM SPS as photo initiator, and the doses test was projected in 1mm glass cuvettes containing the resins. C) Positive and negative resolution tests of the $dECM_{mam}$ and $dECM_{mam}$/Col-I printed with 2000 mJ/cm2 and 1200 mJ/cm2 respectively.

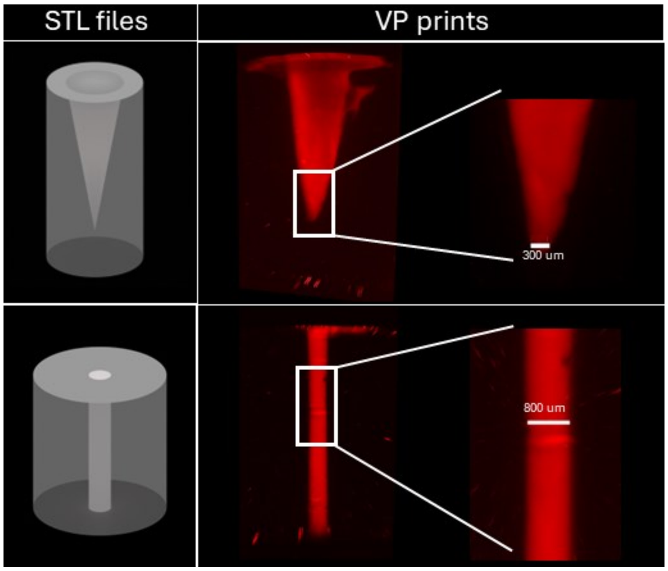


**Figure S17: Volumetric printing: STL files VS. printed constructs**
Constructs printed with $dECM_{mam}$/Col-I (40 mg/ml + 4mg/ml) with 0.5 mM RU and 5mM SPS; 1250 mJ/cm2, printing time: 120 seconds

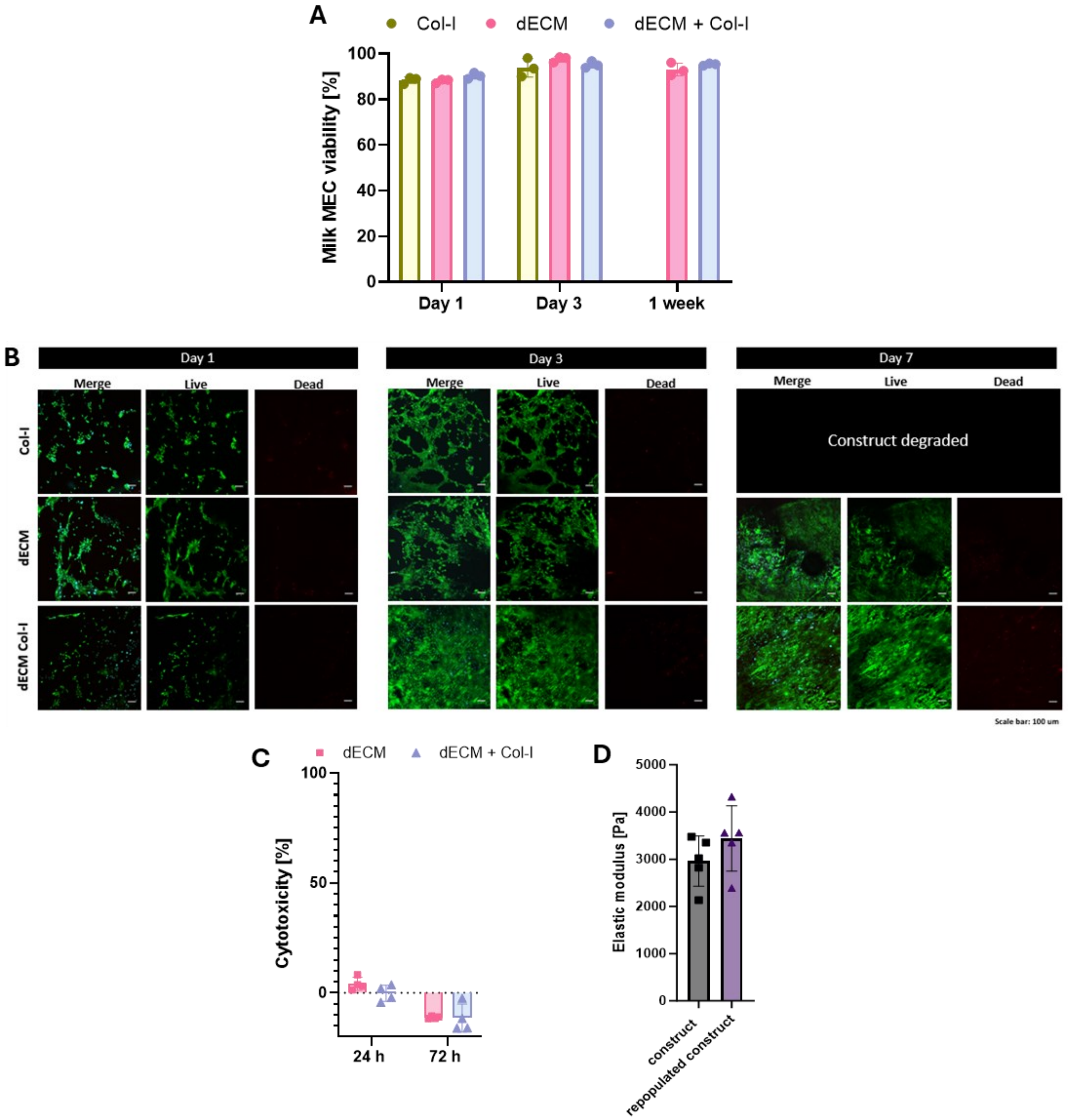


**Figure S18: Biocompatibility of milk mammary epithelial cells (milk MEC) with printed dECM + Col-I constructs.**

A) Milk MEC viability seeded on printed $dECM_{mam}$ and $dECM_{mam}$/Col-I constructs. Viability was assessed with Live/dead staining at day1, 3 and 7 days post seeding (n=3 biological replicates). B) Confocal images of Calcein (green; live) and propidium iodide (red; dead) stained cells seeded on $dECM_{mam}$ and $dECM_{mam}$/Col-I constructs at 1, 3 and 7 days post seeding. C) Bar plot representing the cell cytotoxicity in percent for the milk MEC cultured on the $dECM_{mam}$ and $dECM_{mam}$/Col-I constructs for 24 and 72 hours. Cytotoxicity was assessed using the lactate dehydrogenase assay. D) Elastic moduli of $dECM_{mam}$/Col-I constructs with and without cells seeded after 1 week of culture in 37°C (seeding density: 10.000 cells/cm2). E) Ductal alveolar (200 um) unit repopulated with milk MEC: Measurement of well diameters (n= 3 wells) showing that the shape is maintained over the culture period.

### Full western blot figure 1

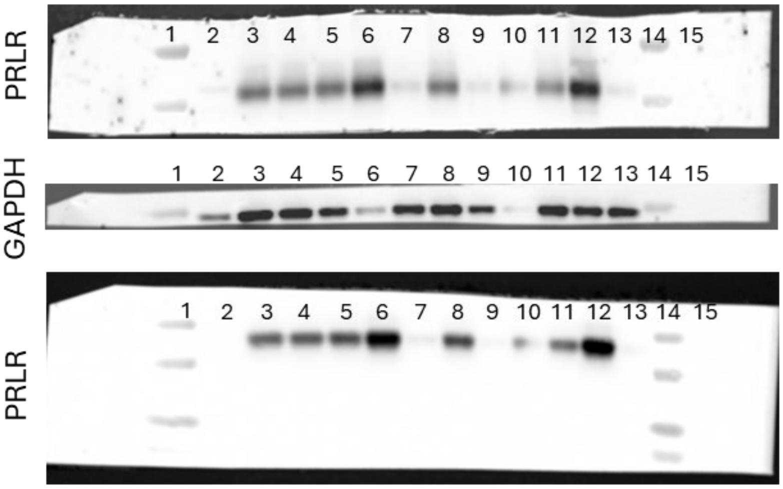


1: ladder
2: Donor 1 P0
3: Donor 1 P1
4: Donor 1 P2
5: Donor 1 P3
6: Donor 2 P0
7: Donor 2 P1
8: Donor 2 P2
9: Donor 2 P3
10: Donor 3 P0
11: Donor 3 P1
12: Donor 3 P2
13:Donor 3 P3
14: ladder
15: empty

### Full western blot figure 5

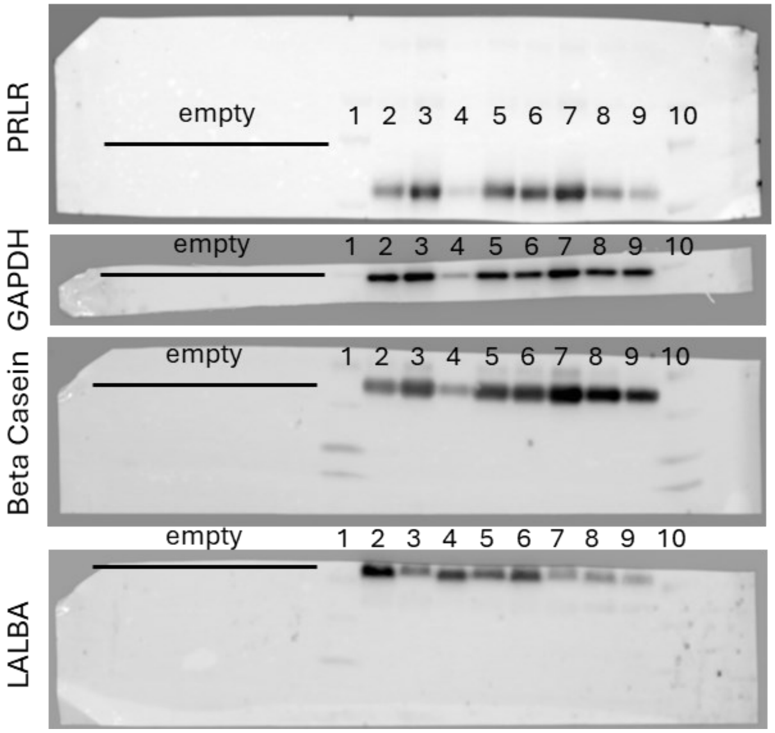


1 & 10: ladder
2: Donor 1 (unstimulated)
3: Donor 2 (unstimulated)
4: Donor 3 (unstimulated)
5: Donor 4 (unstimulated)
6: Donor 1 (100 ng/ml PRL)
7: Donor 2 (100 ng/ml PRL)
8: Donor 3 (100 ng/ml PRL)
9: Donor 4 (100 ng/ml PRL)

**Figure S19: Westernblot full gels**

| Sample | Maternal age [years] | BMI | Infant age [weeks] | Parity | Cells / ml |
|---|---|---|---|---|---|
| Donor 1 | 36 | 23.1 | 35 | 2 | 345K |
| Donor 2 | 27 | 31.7 | 24 | 1 | 734K |
| Donor 3 | 35 | 28.7 | 16 | 1 | 383K |
| Donor 4 | 31 | 21.0 | 21 | 1 | 642 K |

**Table S1. Demographic information of the participating milk donors**

| Antigen | Fluorochrome | Dilution | Clone | Host / isotype | Reactivity | Company | Cat. Number |
|---|---|---|---|---|---|---|---|
| CD90 | BV510 | 1:40 | 5E10 | Mouse IgG1, k | Human | Biolegend | 328125 |
| CD45 | BV785 | 1:80 | HI131 | Mouse IgG1, k | Human | Biolegend | 304008 |
| CD31 | PE | 1:640 | WM59 | Mouse IgG1, k | Human | Biolegend | 303106 |
| CD49f | APC | 1:160 | GoH3 | Rat IgG2a, k | Human | Biolegend | 313615 |
| EPCAM | FITC | 1:160 | 9C4 | Mouse IgG2b, k | Human | Biolegend | 324203 |

**Table S2. List of antibodies used for Flow Cytometry**

| Antigen | Dilution | Host | Reactivity | Company | Cat. Number |
|---|---|---|---|---|---|
| Casein | 1:100 | Mouse | Human | abcam | ab47972 |
| Zonulin1 | 1:100 | Rabbit | Human, MS, Rat, Dog, GP | Invitrogen | 617300 |
| P63 | 1:100 | Mouse | Human | abcam | ab735 |
| CK8 | 1:50 | Rabbit | Human | Invitrogen | PA5118024 |
| CK8 TROMA 1 | 1:50 | Rat | Human, MS | Merck | MABT329 |
| CK14 | 1:100 | Rabbit | Human, MS, Rat | Biolegend | 905304 |
| EPCAM | 1:200 | Rabbit | Human, MS, Rat | BIOSSUSA | bs1513R |
| CD49f | 1:100 | Rat | Human, MS | Invitrogen | 14-0495-82 |
| Milk Fat Globule | 1:100 | Mouse | Human | Neo Biotech. | 4240-MSM1-P0 |
| Col-III | 1:100 | Rabbit | Human | Rockland | 6000401105S |
| Col - I | 1:100 | Mouse | Human | Abcam | ab6308 |

**Table S3. List of primary antibodies used for immunostaining**

| Secondary Antibody | Host | Dilution | Isotype | Company | Cat. Number |
|---|---|---|---|---|---|
| Anti Mouse AF 488 | Goat | 1:500 | IgG (H+L) | Invitrogen | A11001 |
| Anti Rat AF 488 | Goat | 1:500 | IgG (H+L) | Invitrogen | A11006 |
| Anti Rabbit 647 | Goat | 1:500 | IgG (H+L) | Invitrogen | A21244 |
| Anti mouse 568 | Goat | 1:500 | IgG (H+L) | Invitrogen | A11004 |

**Table S4. List of secondary antibodies used for immunostaining**

| Antigen | Dilution | Clone | Host / isotype | Reactivity | Company | Cat. Number |
|---|---|---|---|---|---|---|
| PRLR | 1:1000 | 2E10G5 | Mouse / IgG1 | Human | Proteintech | 67292 |
| PRL | 1:500 | C13 | Mouse | Human | Invitrogen | MA543711 |
| LALBA | 1:1000 | Polyclonal | Rabbit | Human | Invitrogen | PA5-56274 |
| Casein | 1:500 | F20.14 | Mouse | Human | abcam | ab47972 |
| GAPDH | 1:10000 | Polyclonal | Rabbit / IgG | Human | Invitrogen | PA1-987 |

**Table S5. List of antibodies used for Western Blot**

| Gene | Annealing | Oligonucleotide primer sequence |
|---|---|---|
| PRLR | FW | 5'-TTA CCA CAG GGA AGG AGA GAC A-3' |
| | REV | 5'-GGT GTA CTG CTT GCC AAA GTG-3' |
| STAT5A | FW | 5'-GCC GGT TTG AGT GAG GGT TT-3' |
| | REV | 5'-GGA AAC GTG GGA ACA GCA TC-3' |
| ELF5 | FW | 5'-ATT TCC ACT GCA CGC TGG TG-3 |
| | REV | 5'-TGT GTG TCA CCG AGT CCA AC-3' |
| CSN2 | FW | 5'-CCT CTG AGA CTG ATA GTA TTT-3' |
| | REV | 5'-TGG ATG CTG GAG TGA ACT TTA-3' |

**Table S6. List of RT-qPCR primers**
